# Multimodal learning of transcriptomes and text enables interactive single-cell RNA-seq data exploration with natural-language chats

**DOI:** 10.1101/2024.10.15.618501

**Authors:** Moritz Schaefer, Peter Peneder, Daniel Malzl, Mihaela Peycheva, Jake Burton, Anna Hakobyan, Varun Sharma, Thomas Krausgruber, Jörg Menche, Eleni M. Tomazou, Christoph Bock

**Affiliations:** Medical University of Vienna, Institute of Artificial Intelligence, Center for Medical Data Science, Vienna, Austria; CeMM Research Center for Molecular Medicine of the Austrian Academy of Sciences, Vienna, Austria; St. Anna Children’s Cancer Research Institute (CCRI), Vienna, Austria; Doctoral School in Microbiology and Environmental Science, University of Vienna, Vienna, Austria; Max Perutz Labs, Vienna, Austria; Department of Structural and Computational Biology, Center for Molecular Biology, University of Vienna, Vienna, Austria; Ludwig Boltzmann Institute for Network Medicine at the University of Vienna, Vienna, Austria; Faculty of Mathematics, University of Vienna, Vienna, Austria; Medical University of Vienna, Center for Cancer Research, Vienna, Austria

**Keywords:** Single-cell RNA-seq analysis, multimodal deep learning, large language models, bioinformatics

## Abstract

Single-cell RNA-seq characterizes biological samples at unprecedented scale and detail, but data interpretation remains challenging. Here we introduce CellWhisperer, a multimodal machine learning model and software that connects transcriptomes and text for interactive single-cell RNA-seq data analysis. CellWhisperer enables the chat-based interrogation of transcriptome data in English language. To train our model, we created an AI-curated dataset with over a million pairs of RNA-seq profiles and matched textual annotations across a broad range of human biology, and we established a multimodal embedding of matched transcriptomes and text using contrastive learning. Our model enables free-text search and annotation of transcriptome datasets by cell types, states, and other properties in a zero-shot manner and without the need for reference datasets. Moreover, CellWhisperer answers questions about cells and genes in natural-language chats, using a biologically fluent large language model that we fine-tuned to analyze bulk and single-cell transcriptome data across various biological applications. We integrated CellWhisperer with the widely used CELLxGENE browser, allowing users to interactively explore RNA-seq data through an integrated graphical and chat interface. Our method demonstrates a new way of working with transcriptome data, leveraging the power of natural language for single-cell data analysis and establishing an important building block for future AI-based bioinformatics research assistants.

## Introduction

Transcriptome profiling is widely used for the characterization of cells and tissues (Moreno et al. 2022; Quake 2021). Bulk RNA sequencing (RNA-seq) provides a detailed assessment of cell states and biological functions through a single cost-effective assay (Stark et al. 2019), and with single-cell RNA sequencing (scRNA-seq), researchers can disentangle the cell composition and the biological heterogeneity of tissues, organs, and diseases (Aldridge and Teichmann 2020). This technology has enabled global efforts to establish genome-wide profiles for all cell types in the human body (Regev et al. 2018), working toward a molecular atlas of human physiology and a reference map for investigating regulatory changes associated with human diseases.

A typical scRNA-seq dataset can be represented by a count matrix with ∼20,000 genes (columns) and thousands or millions of single cells (rows). Analyzing and interpreting such datasets is a complex task that requires both bioinformatic skills and application-specific biological expertise. To facilitate scRNA-seq data analysis, software tools have been developed for a wide range of tasks including data visualization, cell clustering, cell type annotation, differential expression, and gene set analysis (Zappia and Theis 2021). Moreover, deep learning-based “foundation models” trained on large scRNA-seq datasets promise to go beyond specialized tools and support analysis tasks they were not explicitly optimized for (Simon et al. 2024; Szałata et al. 2024).

Here we demonstrate scRNA-seq data exploration with natural language, allowing the user to interrogate cells in English language, with no need to adhere to any particular format or syntax. Our CellWhisperer model and software supports free-text search (e.g., *‘Show me tissue-resident T cells in the intestine’*) and answers a broad range of questions about cells (e.g., *‘What are these selected cells?’, ‘Which genes are highly expressed in these cells?’, ‘What is the role of KLRD1 in NK cells?’*). The model’s response is based on the combination of the scRNA-seq data and the biological knowledge of a large language model (LLM), resulting in answers such as: ‘*The selected cells appear to be CD16+ NK cells, which are a subset of natural killer (NK) cells that play a crucial role in the innate immune response […]’, ‘The top expressed genes in these cells include NKG7, KLRD1, GNLY, GZMA, PRF1 […]’, ‘KLRD1 (CD94) is a receptor that plays a role in NK cell activation and cytotoxicity. It can recognize MHC class I molecules on target cells and trigger NK cell-mediated cytotoxicity’*.

We trained CellWhisperer to integrate RNA profiles and their metadata annotations by multimodal contrastive learning (Radford et al. 2021), creating a joint multimodal embedding of transcriptomes and text. The training data comprised over a million transcriptomes and their natural-language annotations, which we prepared with AI-assisted curation from two large community-wide repositories: Gene Expression Omnibus (GEO) (Clough et al. 2024; Edgar et al. 2002) and CELLxGENE Census (CZI Single-Cell Biology Program et al. 2023). To enable interactive chat-based data exploration, CellWhisperer incorporates an open-source LLM (Jiang et al. 2023; H. Liu et al. 2023) that we fine-tuned to answer questions about cell states while considering user-provided transcriptome profiles. For user-friendly analysis of scRNA-seq data, we integrated CellWhisperer into the widely used CELLxGENE Explorer (Megill et al. 2021). The CellWhisperer software is available at https://cellwhisperer.bocklab.org, and usage examples are provided in **Fig. 3** and **Supplementary Video 1**.

In summary, we developed an AI interface that establishes natural language as an intuitive channel for interacting with scRNA-seq datasets, enabled by multimodal integration of transcriptomes and text, and exploiting the broad biological knowledge that is embedded into CellWhisperer’s multimodal LLM. We envision such natural-language interactions with data as a key element of future AI-based bioinformatics research assistants.

## Results

### CellWhisperer links transcriptomes and text through a multimodal embedding model and LLM

To enable scRNA-seq data analysis using natural language, we created CellWhisperer, a multimodal AI model of transcriptomes and text that supports data-centric conversations about cells and genes. CellWhisperer was built in three steps (**Fig. 1a**): (i) LLM-assisted curation of training data, resulting in over a million pairs of human RNA-seq profiles and matched textual annotations; (ii) learning of a multimodal embedding model that connects text to transcriptomes (e.g., cell search using free-text queries) and transcriptomes to text (e.g., cell annotation with textual descriptions); (iii) transcriptome-aware fine-tuning of an LLM for data-centric question answering and natural-language chats. This section describes each of these three steps, while further technical details are provided in the **Methods** section, in **Supplementary Note 1** and **2**, and in **Extended Data Fig. 1**.

**Figure 1.**
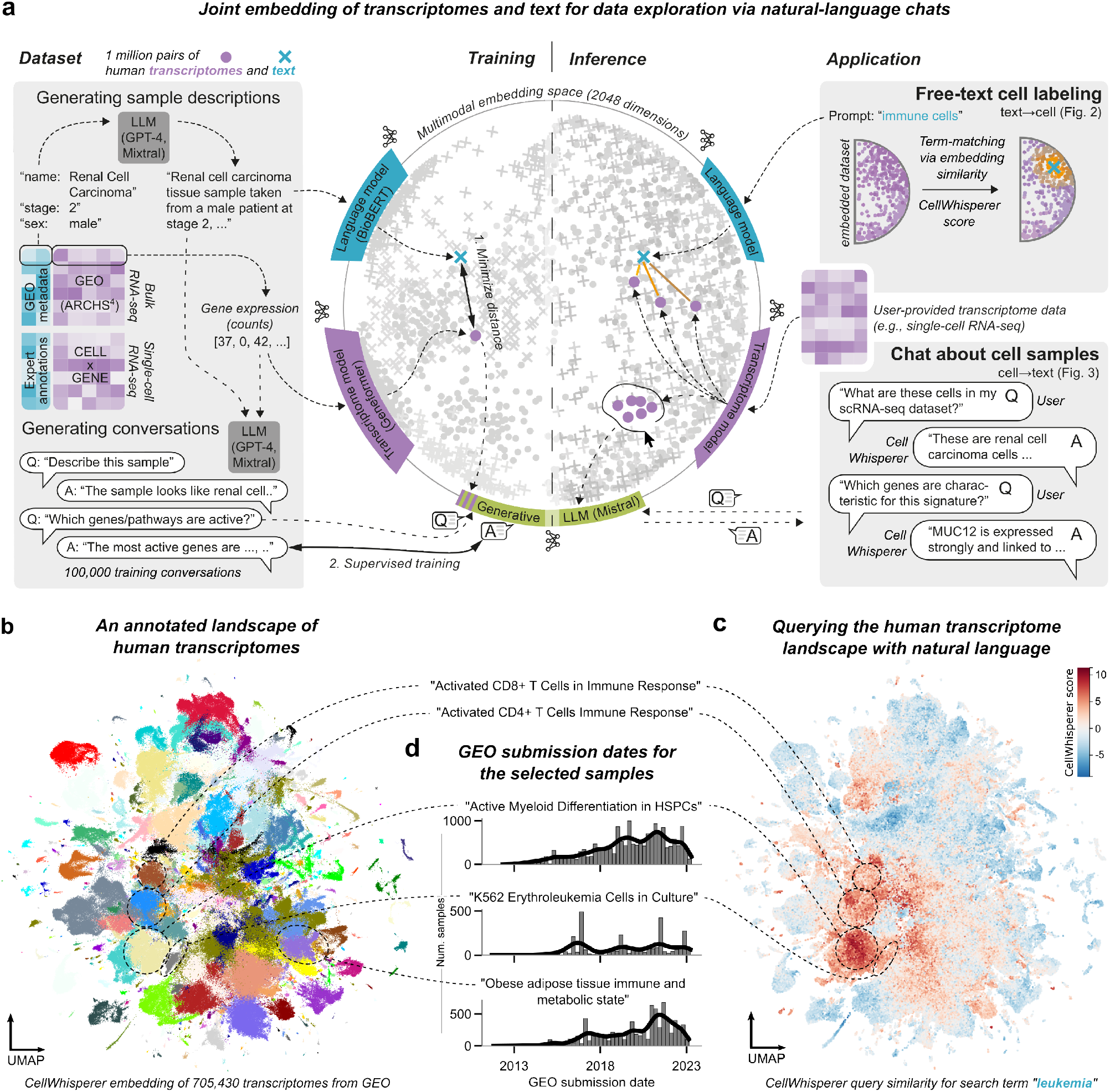
Overview of the CellWhisperer multimodal AI for natural-language analysis of transcriptome data. a) Conceptual overview of training dataset generation (left), model training and inference (center), and applications in scRNA-seq data analysis (right). b) CellWhisperer embeddings for human transcriptomes from the GEO repository visualized as a UMAP. Clusters were computed using the Leiden algorithm, and cluster labels were generated by CellWhisperer based on cluster-averaged transcriptome embeddings. The fully annotated dataset is available for interactive browsing and analysis on the project website (https://cellwhisperer.bocklab.org/geo). c) CellWhisperer scores for the free-text query term ‘leukemia’ projected on the human transcriptome landscape (from panel b). d) Retrieval of sample metadata (here: GEO submission date) for transcriptomes interactively selected in CellWhisperer.

First, we created a training dataset that connects transcriptomes (i.e., RNA profiles for bulk samples or single-cell clusters) with concise textual annotations (e.g., *‘Renal cell carcinoma tissue sample taken from a male patient at stage 2, with no metastasis, preserved in FFPE blocks’*), across a broad range of cell types and conditions. To that end, we performed LLM-assisted curation of the Gene Expression Omnibus (GEO) and CELLxGENE Census data repositories. GEO comprises RNA-seq data from ∼20,000 individual studies based on researcher submissions. Its inherent diversity is a great advantage, as it covers essentially all fields of human biology that have been studied with RNA-seq; but it also creates the need for data harmonization. To obtain coherent transcriptome profiles, we built upon the ARCHS^4^ project, which carried out uniform processing of RNA-seq data from GEO (Lachmann et al. 2018); and to retrieve concise and biologically descriptive textual annotations for each sample, we developed an LLM-assisted curation procedure based on sample-associated metadata (which comprise semi-structured information such as cell types, organs, tissues, and diseases, description of experimental methods, scientific project abstracts, etc.). This way, we derived from GEO a coherent training dataset of 705,430 human transcriptomes with matched textual annotations.

Our second data source, CELLxGENE Census, comprises several hundred scRNA-seq datasets including reference maps from the Human Cell Atlas. We grouped the cells in each dataset based on the provided metadata and generated pseudo-bulk transcriptomes by averaging across the scRNA-seq profiles within each group. We then applied our LLM-assisted curation procedure to condense the metadata for each group into concise biological descriptions, resulting in 376,983 human transcriptomes with matched textual annotations.

Second, we used the combined dataset with 1,082,413 pairs of transcriptomes and their textual annotations to train a deep learning model that integrates the two modalities into a joint multimodal embedding space. We adapted the contrastive language-image pretraining (CLIP) architecture (Radford et al. 2021) and processed the transcriptome profiles with the Geneformer model for gene expression (Theodoris et al. 2023), while the textual annotations were processed with the BioBERT model for biomedical text (Lee et al. 2020). Each of the two resulting vectors was mapped into a 2048-dimensional multimodal embedding space using conventional feed-forward neural network layers. These adapter layers and the BioBERT model were trained to place the two modality-specific embeddings within close proximity in the joint embedding space. We validated that the resulting CellWhisperer multimodal embedding was capable of retrieving the transcriptome corresponding to a given textual annotation and vice versa **–** a standard metric of CLIP model performance (Radford et al. 2021) **–** and we obtained a mean ROC-AUC value of 0.927 (**Extended Data Fig. 1a**).

The trained CellWhisperer model can be prompted with free-text queries to search for transcriptomes that match the search query. For example, the *‘immune cells’* query is processed with the BioBERT-based language model, the resulting embedding is compared to the embeddings of all relevant cells or samples from the Geneformer-based transcriptome model, resulting in a quantitative measure (the *‘CellWhisperer score’*) that assesses the match between the query and a given transcriptome. High CellWhisperer scores thus indicate that the model will rank this transcriptome highly in response to this free-text query and report it with high priority.

Third, to enable natural-language chats with CellWhisperer, we customized and fine-tuned the Mistral 7B open-source LLM (Jiang et al. 2023) to incorporate transcriptome information in addition to textual input. Our approach is inspired by the ability of multimodal LLMs – such as GPT-4, Gemini, and the open-source LLaVA model (H. Liu et al. 2023) – to interpret and converse about images. We generated a training dataset of 106,610 question-answer-style conversations, including simple rule-based conversations (e.g., *‘What does the sample represent?’*, with the sample’s textual annotation as the designated answer) as well as more complex LLM-generated conversations that all take transcriptome profiles into account (see **Methods** section for technical details and **Supplementary Note 1** for examples). Based on this training dataset, we used the CellWhispererderived transcriptome embeddings together with the prepared questions as input to the Mistral 7B LLM (with an adapter layer that converts the transcriptome embeddings into *Mistral*-compatible token-level embeddings), and we fine-tuned this LLM to produce the matched answers. The resulting fine-tuned LLM responds to freetext questions and engages in natural-language chats about cells and their biological functions, gene-regulatory mechanisms, and other biological processes that can be linked to transcriptional cell states.

To illustrate CellWhisperer’s multimodal embedding model and the fine-tuned conversational LLM, we clustered the CellWhisperer embedding vectors for the 705,430 GEO-derived human transcriptomes and annotated each cluster with a descriptive text that CellWhisperer generated from the cluster-averaged transcriptomes (see **Fig. 1b** for a UMAP representation and https://cellwhisperer.bocklab.org/geo for an interactive version). The CellWhisperer embedding successfully captured cell types, developmental stages, tissues, diseases, and more. We queried the multimodal model with search term *‘leukemia’* and projected the CellWhisperer score (i.e., the match between the query and each transcriptome) on top of the UMAP (**Fig. 1c**). This query finds biologically fitting clusters (e.g., *‘Active Myeloid Differentiation in HSPCs’*). As each data point connects back to an actual sample in the GEO database, we can retrieve the corresponding sample metadata and for example assess the popularity of RNA-seq analysis for certain cell clusters and biological functions over the last decade (**Fig. 1d**).

In summary, we present a multimodal AI that facilitates the seamless transition from transcriptomes to text and vice versa, which enables the chat-based analysis of bulk and scRNA-seq data in English language.

### CellWhisperer captures and describes diverse cell properties via zero-shot classification and text generation

To assess how well CellWhisperer’s multimodal embedding model has learned relevant aspects of human biology, we tested its ability to predict cell types, diseases, tissues, and organs in a zero-shot manner (i.e., without any task-specific training or reference data). To that end, we used well-annotated transcriptome datasets excluded from CellWhisperer’s training data, and queried CellWhisperer with text about cell properties (e.g., cell type *‘erythrocytes’*) to identify the corresponding cells (**Fig. 2a**). We then calculated the coherence between the CellWhisperer predictions and the original dataset annotations (which we treated as ground truth).

**Figure 2.**
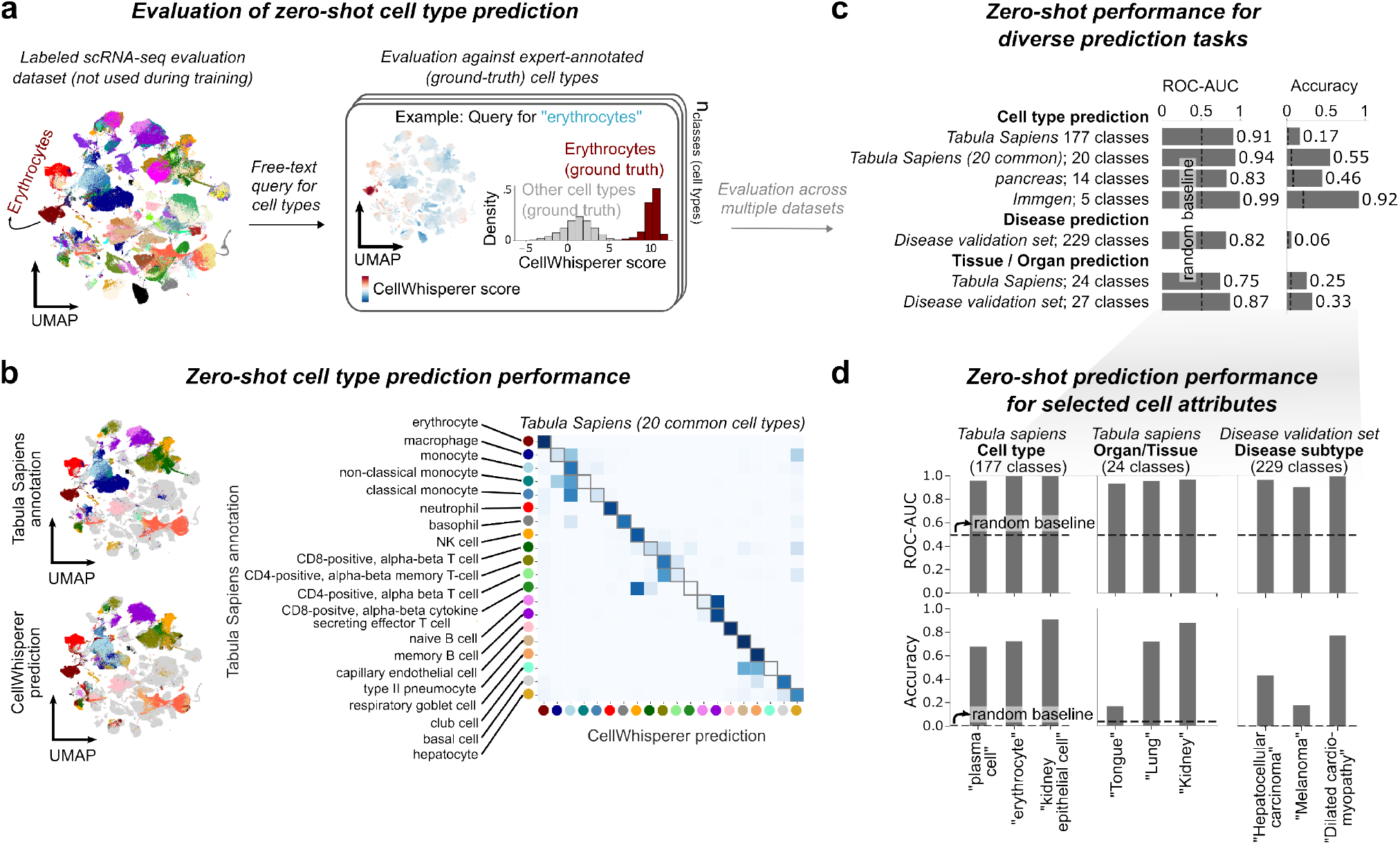
Evaluation of the CellWhisperer model by zero-shot prediction of cell types, diseases, and tissues. a) Outline of zero-shot performance evaluation for the CellWhisperer multimodal model. Left: UMAP representation of CellWhisperer transcriptome embeddings for the Tabula Sapiens dataset, colored by the dataset’s expert-annotated cell types used as “ground truth”. Right: CellWhisperer scores for the free-text query ‘erythrocytes’ projected on the UMAP and as histograms grouped by the “ground truth” erythrocyte label. b) Comparison of “ground truth” cell types (top UMAP) and CellWhisperer predictions (bottom UMAP) for the Tabula Sapiens (20 common cell types) dataset, with the confusion matrix as a heatmap (right). c) Zero-shot performance for diverse prediction tasks (cell type, disease subtype, tissue/organ of origin), calculated across multiple single-cell and bulk RNA-seq datasets and quantified by macro-averaged ROC-AUC and accuracy values. Dotted black lines denote random baseline performances corresponding to 0.5 (for ROC-AUC) and 1/number of classes (for accuracy). d) Examples of CellWhisperer zero-shot prediction performance for selected tasks (from panel c).

For the *Tabula Sapiens dataset*, which comprises scRNA-seq profiles for 483,152 cells from 24 organs (Tabula Sapiens Consortium et al. 2022), CellWhisperer distinguished 20 frequent cell types with a ROC-AUC value of 0.94 (**Fig. 2b-c**). Deviations were for the most part between closely related cell types, for example between cells annotated as “monocytes” versus “classical monocytes” and between subgroups of T cells (**Fig. 2b**). We also tested the prediction performance for the entire set of 177 annotated cell types in the *Tabula Sapiens dataset* and obtained a ROC-AUC value of 0.91, but with a lower accuracy value given the large number of classes with many highly similar cell types (**Fig. 2c**). Moreover, for bulk RNA-seq profiles of immune cells from the Immgen consortium (Heng et al. 2008; Immunological Genome Project Consortium 2023), we obtained a ROC-AUC value of 0.99, and for a challenging scRNA-seq dataset of human pancreas with closely related cell types and pronounced batch effects (Luecken et al. 2022), the ROC-AUC value was 0.83 (**Fig. 2c**).

Notably, CellWhisperer was never specifically trained to predict cell types. Rather, its ability to link cell type labels to typical scRNA-seq profiles of such cells emerged from the learned connection between transcriptomes and their textual annotations (which tend to include information about cell types). We expanded this zero-shot analysis to test whether CellWhisperer can also predict other information contained in metadata-derived textual annotations, such as diseases, tissues, and organs. To that end, we assembled a collection of 14,112 disease-associated transcriptomes from GEO, which we excluded from the training data. Predicting 229 disease subtypes represented in this *disease validation dataset*, CellWhisperer achieved a ROC-AUC value of 0.82 (**Fig. 2c**), indicating that disease prediction is harder than cell type prediction but possible with a performance that is substantially better than random guessing. Similarly, CellWhisperer was able to predict the tissue-of-origin of bulk and single-cell transcriptomes with better-than-random prediction performance both in the *Tabula Sapiens dataset* (ROC-AUC: 0.75) and in the *disease validation dataset* (ROC-AUC: 0.87) (**Fig. 2c, d**).

To gauge the breadth of biological processes captured by our model, we investigated its recognition of expert-curated gene sets spanning diverse areas of biology. For each of 8,812 gene sets, we used the gene set label (e.g., *‘colorectal cancer’*) as a query text to CellWhisperer and determined how well each sample in our *disease validation dataset* matched the query. We then calculated the correlation between this pure text-based assessment (which does not use the information of which genes are part of the gene set) and the gene expression enrichment across all genes in the gene set, for each sample in the *disease validation dataset* (**Extended Data Fig. 2a**). In other words, we tested if CellWhisperer had learned an implicit understanding of genes that matter for a given biological concept, based on the gene set label. We indeed found a clear positive association between CellWhisperer scores for gene set labels and the expression of the corresponding gene sets (**Extended Data Fig. 2b-c, Supplementary Table 1**), indicating that our model has implicitly learned (albeit imperfectly) many of the tested biological concepts. Importantly, CellWhisperer achieved this by training on transcriptomes and their textual representations, without having seen any expert-curated gene sets during model training.

For further evaluation, we tested how well the CellWhisperer model can distinguish between biological signal and technical noise, based on an established batch effect removal benchmark (Luecken et al. 2022). We observed improved performance of CellWhisperer compared to the transcriptome-only Geneformer model (**Extended Data Fig. 2d**), suggesting that incorporation of textual information, as done in CellWhisperer’s training, not only enables new functionality but may also increase the quality of the transcriptome embedding.

Finally, we benchmarked CellWhisperer’s multimodal LLM using the perplexity metric (Rosenfeld 2000), a common evaluation criterion for LLMs. First, we assessed how well each question-answer pair fits with the matched transcriptome. We observed a 90% preference for matched over unmatched transcriptomes in our *conversation evaluation dataset* with 200 question-answer pairs (**Extended Data Fig. 3a**), confirming that our LLM meaningfully incorporated the transcriptome embedding into the response generation. Second, we found that most cell type labels showed a preferential association with transcriptomes of the corresponding cell type in the *Tabula Sapiens dataset (20 common cell types)* (**Extended Data Fig. 3b**).

In conclusion, multiple lines of evidence (zero-shot prediction of cell types, diseases, tissues, and organs, a batch effect removal task, gene set prediction from their labels, perplexity for question-answer pairs, and perplexity for cell type labels) support our hypothesis that CellWhisperer has learned a meaningful model of cells and relevant biological processes, solely based on training data of matched transcriptomes and text.

### CellWhisperer enables chat-based analysis of scRNA-seq data through a user-friendly web-based interface

To make CellWhisperer broadly accessible for chat-based analysis of transcriptome data, we integrated it into CELLxGENE Explorer through a dedicated chat box (**Fig. 3a**; https://cellwhisperer.bocklab.org; **Supplementary Video 1**). CELLxGENE Explorer is a web-based tool and graphical user interface for analyzing scRNAseq profiles. It supports the visual inspection, selection, and filtering of cells and annotations, as well as the identification of differentially expressed genes. Complementing the core functionality of CELLxGENE Explorer, the CellWhisperer chat box enables data exploration through natural language, such as free-text search for cells with a wide range of properties, automatic annotation of cell clusters, chat-based investigation of interactively selected cells, and more generally the discussion of cells and genes in natural language.

**Figure 3.**
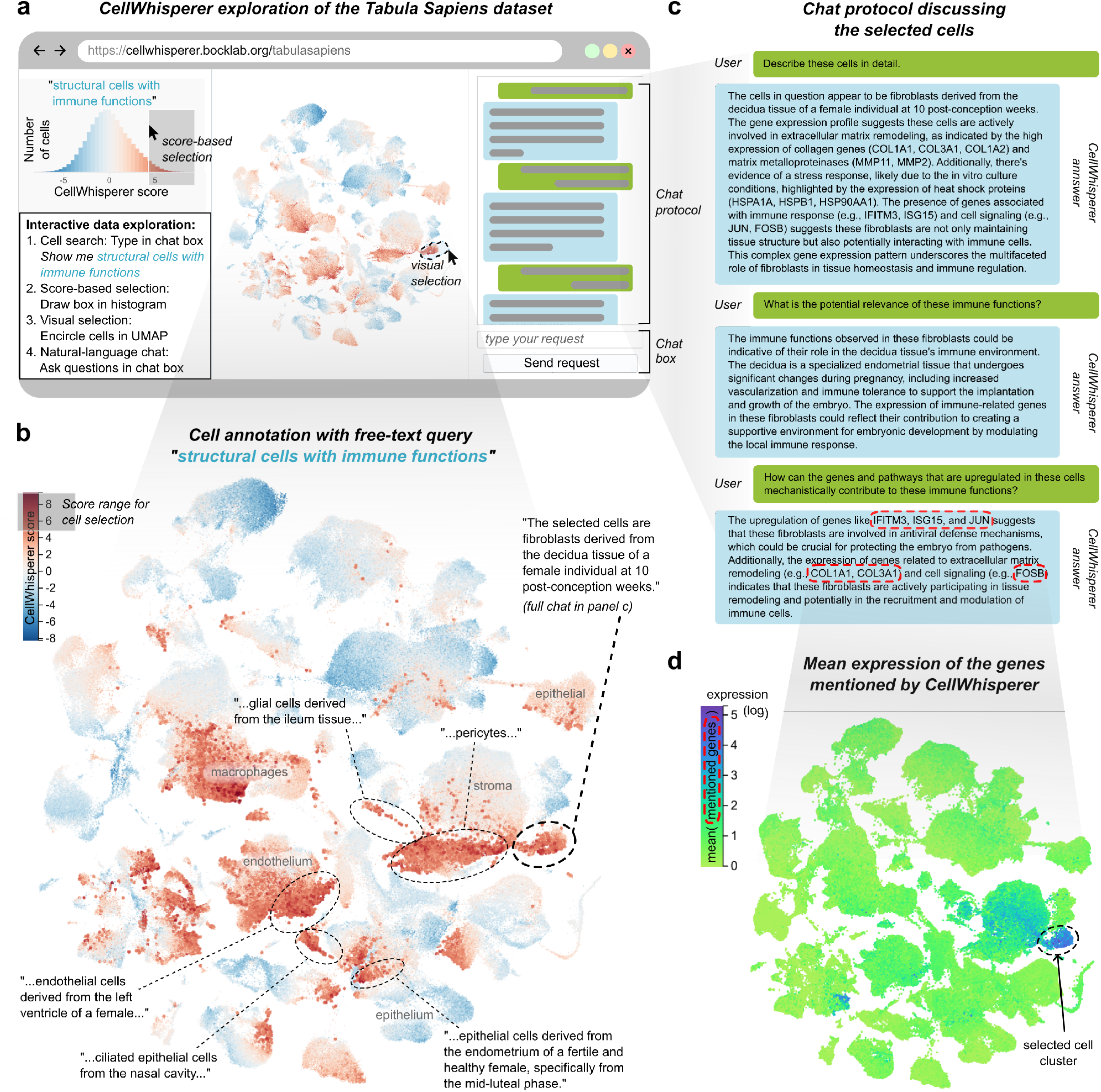
Interactive chat-based exploration of scRNA-seq data using CellWhisperer. a) Schematized screenshot of the CellWhisperer web application, showing the Tabula Sapiens dataset with CellWhisperer scores for the free-text query ‘Show me structural cells with immune functions’. b) Zoom-in on the UMAP of CellWhisperer embeddings and scores (from panel a). Clusters of cells with high CellWhisperer scores for the free-text query were interactively selected in the web application and labeled by prompting CellWhisperer’s multimodal LLM for a natural-language description (chat request: ‘Describe the selected cells’). Shown responses were trimmed to the most relevant parts, as indicated by ‘…’. Annotations in grey font were added manually to the CellWhisperer plot. c) Screenshot of a CellWhisperer conversation about the interactively selected cells (marked in panel b). d) Mean-aggregated expression of the genes IFITM3, ISG15, JUN, COL1A1, COL3A1, and FOSB, which were mentioned in the CellWhisperer responses (from panel c), interactively projected on the Tabula Sapiens UMAP using the ‘Gene Sets’ feature in the CellWhisperer web application.

We illustrate CellWhisperer’s use and utility for scRNA-seq analysis on the well-characterized *Tabula Sapiens* dataset (**Fig. 3**). In previous work, we described widespread immune gene activity in non-hematopoietic “structural cells” of the mouse (Krausgruber et al. 2020), prompting us to explore this phenomenon in a large multiorgan human scRNA-seq dataset. We thus entered *‘structural cells with immune functions’* into the CellWhisperer chat box and obtained the corresponding CellWhisperer score as a color-coded overlay to the UMAP representation of the *Tabula Sapiens* dataset (**Fig. 3a,b**). Among the cells that scored high for this query were endothelial cells, epithelial cells, fibroblasts, and pericytes (**Fig. 3b**), which are all known or suspected to play important immune-regulatory roles (Amersfoort et al. 2022; Davidson et al. 2021; Larsen et al. 2020).

To investigate these cells in more detail, we sequentially selected cell clusters with high CellWhisperer scores (by drawing a circle around the cells of interest) and prompted CellWhisperer with *‘Describe these cells in detail’* through the chat box (**Fig. 3a-c**). We obtained textual descriptions for each cell cluster, which were generated by the CellWhisperer model based on the average transcriptome across the selected cells as well as the query (**Fig. 3b**). The resulting descriptions often contained information about cell types, organs, and developmental stages, and occasionally information about potential sample donors (e.g., male or female), highly expressed genes (e.g., genes encoding collagens and matrix metalloproteinases in fibroblasts), biological functions (e.g., stress response), and other details. We observed that the generated descriptions often referred to potential immune functions of the selected cells, consistent with our initial search query.

To obtain additional information about these cells, we interactively selected one of the cell clusters and asked two follow-up questions (*‘What is the potential relevance of these immune functions?’* and *‘How can the genes and pathways that are upregulated in these cells mechanistically contribute to these immune functions?’*) resulting in a coherent conversation with CellWhisperer, which provided further characterization in the form of highlighted genes and biological functions that are relevant in the selected cells (**Fig. 3c**). As a plausibility check, we confirmed the expression of those genes by mapping them on the UMAP representation (**Fig. 3d**).

Taken together, the CellWhisperer chat box – integrated into the popular CELLxGENE Explorer – provides user-friendly access to CellWhisperer’s AI features and demonstrates the complementarity of visual inspection and natural-language chats for the interactive exploration of scRNA-seq data.

### CellWhisperer enables interactive explorative analysis of user-provided transcriptome datasets

To incorporate user-provided transcriptome datasets into CellWhisperer, we built a data processing pipeline that computes CellWhisperer embeddings and clustered UMAPs with CellWhisperer-generated captions, starting from scRNA-seq read count matrices. Technical instructions and the source code for data processing and running the CellWhisperer web application are provided at https://github.com/epigen/cellwhisperer. The results of this data processing are saved in a single file for convenient storage and import into CellWhisperer, while also facilitating reproducibility and sharing of CellWhisperer analyses. Here we illustrate the analysis of user-provided data, investigating stem/progenitor cells in the colon and their response to inflammatory environments, based on a published scRNA-seq read count matrix of pathogenic and adjacent normal biopsies from patients with inflammatory bowel disease and healthy controls (Parikh et al. 2019) (**Fig. 4a**).

**Figure 4.**
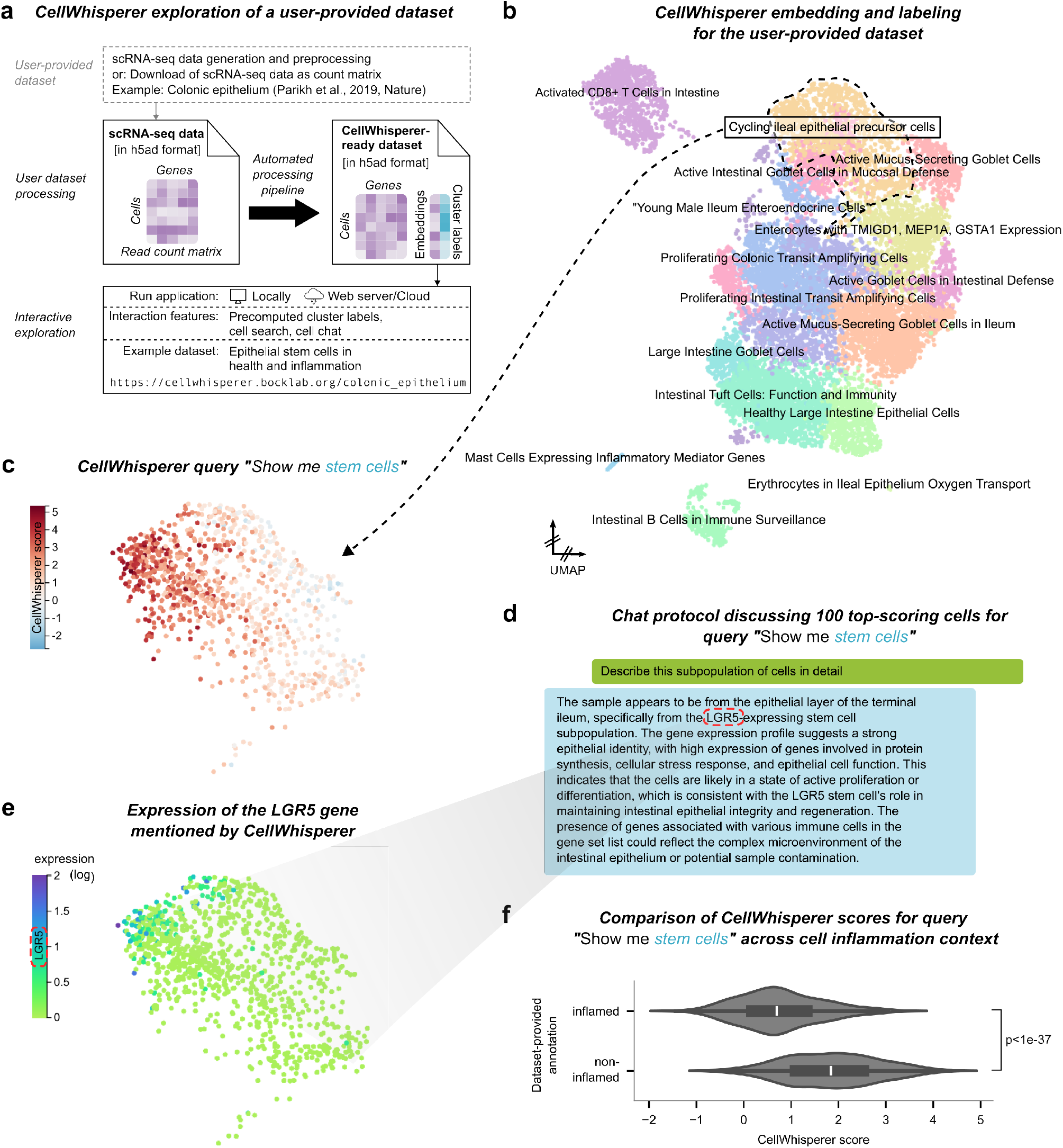
Interactive exploration of user-provided scRNA-seq data with CellWhisperer. a) Outline of the preprocessing and import of user-provided scRNA-seq datasets into CellWhisperer. b) UMAP of the colonic epithelium dataset imported into CellWhisperer. This dataset comprises scRNAseq profiles of inflamed and non-inflamed colonic epithelium from patients with inflammatory bowel syndrome as well as healthy controls. Cluster labels shown on the UMAP were generated by Cell- Whisperer, and clusters were repositioned for compactness. The dataset can be explored interactively on the CellWhisperer website (https://cellwhisperer.bocklab.org/colonic_epithelium). c) Zoom-in on the cluster labeled as ‘Cycling ileal epithelial precursor cells’, colored by CellWhisperer scores for the free-text query: ‘Show me stem cells’. d) CellWhisperer chat about the top 100 cells with highest CellWhisperer scores for the ‘Show me stem cells’ query (from panel c). e) Expression levels of the LGR5 gene mentioned by CellWhisperer (in panel d), plotted for the cell cluster labeled as ‘Cycling ileal epithelial precursor cells’. f) Comparison of CellWhisperer scores for the ‘Show me stem cells’ query (from panel c) between cells that were annotated as inflamed in the colonic epithelium dataset (i.e., sampled from inflamed lesions) or as non-inflamed, across all cells in the cluster annotated as ‘Cycling ileal epithelial precursor cells’. Standard violin plot are shown with inner boxplots corresponding to the interquartile range and whiskers extending to the farthest data point within 1.5 times the interquartile range. Distributions were compared using the two-sample Kolmogorov-Smirnov test.

The CellWhisperer-generated cell cluster annotations provide an overview of this dataset and its contributing cell types. These comprise various types of epithelial cells, such as *‘Cycling ileal epithelial precursor cells’* and *‘Large Intestine Goblet Cells’*, as well as immune cells, including *‘Activated CD8*^*+*^ *T Cells in Intestine’* and *‘Mast Cells Expressing Inflammatory Marker Genes’* (**Fig. 4b**). Focusing on the *‘Cycling ileal epithelial precursor cells’*, we searched for cells with stem cell characteristics using the CellWhisperer query *‘Show me stem cells’* and identified cells within this cluster that scored highly for this query (**Fig. 4c**). Further investigation of these putative stem cells in a series of additional questions and answers with CellWhisperer (**Fig. 4d**) suggested that a subset of these cells are *LGR5*-expressing epithelial stem cells, which constitute well-established stem cells of the gut (Clevers 2013). As expected, *LGR5* gene expression (**Fig. 4e**) was highly correlated with the CellWhisperer score for the *‘Show me stem cells’* query (**Fig. 4c**).

Finally, the observed biological heterogeneity of the *‘Cycling ileal epithelial precursor cells’* cluster (**Fig. 4c, e**) prompted us to compare the prevalence of the CellWhisperer-annotated epithelial stem cells between inflamed and non-inflamed colon samples. We found that the non-inflamed samples were characterized by more pronounced CellWhisperer scores for *‘stem cells’* compared to the inflamed samples (**Fig. 4f**). This analysis suggests that chronic gut inflammation in patients with inflammatory bowel disease negatively impacts *LGR5*- expressing epithelial stem cells, which matches conclusions of the study from which the dataset was obtained (Parikh et al. 2019) as well as previous *in vitro* results (Wang et al. 2019).

Importantly, all these analyses were performed interactively using CellWhisperer integrated into CELLxGENE Explorer. Overall, we thus provide a user-friendly and flexible approach for custom scRNA-seq data exploration – driven by the synergistic nature of visual inspection and chat-based analysis using natural language.

## Discussion

Transcriptome profiling is widely used for characterizing the biological state of cells and tissues, but data analysis and biological interpretation remain challenging. Here we demonstrate scRNA-seq data exploration with natural language, using a multimodal AI model that combines transcriptome profiles with an understanding of biological text, and an LLM chat interface for interactive investigation of cell states. The performance observed in our evaluations suggests that multimodal models of transcriptomes and text can already facilitate exploratory analysis of biomedical data, and natural language may evolve into a widely used additional channel for interactive analysis of biomedical data, complementing visual data inspection and programming-based data analytics. We also envision that natural language may become an attractive, human-interpretable integration layer by which AI models of different scales (e.g., foundation models of molecules, cells, organs, and patients) will interact and exchange their perspectives on a shared question, thereby facilitating multiscale data analysis.

Natural language and chat-based user interfaces hold great potential to facilitate and enhance biological data analysis. As illustrated by the CellWhisperer integration into CELLxGENE Explorer, chat-based analysis will often complement rather than replace visual approaches, thus adding a new channel to the interactive analysis of biological datasets. This has the potential to make data exploration more fluid and creative, as the user is unburdened by the need to memorize complex syntax or look up genes, and there is always the option to ask follow-up questions for clarification. It also reduces barriers to entry, for example for biologists with no programming experience and a natural preference to interact in English language. Moreover, by connecting the chat functionality to voice recognition, it may become possible to interact verbally with the AI, for example in the context of virtual reality data analysis software or for researchers with vision impairment; and given the multi-language capabilities of many LLMs, it will be feasible to support languages other than English. Data analysis in natural language may thus contribute to making bioinformatics more inclusive in several directions.

CellWhisperer builds on several recent advances in AI methodology. First, to establish our coherently annotated, community-scale training dataset of over a million bulk and pseudo-bulk transcriptomes, we used general-purpose LLMs for AI-assisted data curation. Second, CellWhisperer employs powerful modality-specific embedding models to process its inputs of transcriptomes (with Geneformer) and text (with BioBERT). Third, we adapted elements of the CLIP and LiT architectures for learning multimodal embeddings of transcriptomes and matched textual annotations, which constitutes the foundation of the CellWhisperer model and software. Fourth, inspired by image-recognizing chat bots and other multimodal applications of LLMs (Yin et al. 2023), we modified a general-purpose LLM to support chat-based analysis of scRNA-seq data by fine-tuning it with 106,610 AI-generated transcriptome-centric conversations about cells and biological processes.

The current version of CellWhisperer constitutes a proof-of-concept that is useful for interactive exploration of scRNA-seq data, with the following caveats. First, like other LLMs, CellWhisperer does not understand the user questions and its own responses in a human sense; rather, it has learned to continue the conversation based on large amounts of training data on how transcriptome-centric question-answering usually unfolds. CellWhisperer should thus be considered a tool for exploratory data analysis; it should not be trusted blindly and without validation. Second, CellWhisperer relies on domain-specific models for its embedding of transcriptomes and text, and on an LLM for text generation. It thus inherits some of their limitations. For example, Geneformer cannot currently process more than 2048 genes per cell and lacks quantitative resolution. To let CellWhisperer profit from progress with these domain-specific models, we implemented a highly modular software architecture that makes it easy to swap the underlying models. Third, CellWhisperer’s LLM occasionally “hallucinates”, most frequently by providing overly specific information about the sample origin (e.g., *‘T cells from a 85 year old male’*). This is the result of the abundance of such information in the training data and could possibly be addressed by further data curation or by fine-tuning with human feedback for the chat responses. Fourth, CellWhisperer can only be as good as the available training data, and areas of biology that are not well represented in public databases are unlikely to be modeled well by CellWhisperer or similar models.

We also performed a risk assessment of the CellWhisperer method and software, given concerns about inherent risks of modern AI (Bengio et al. 2024). We concluded that CellWhisperer can be considered low-risk, enabling us to make all aspects of the technology openly accessible to the general public. We identified as the most relevant risk of CellWhisperer that incorrect answers may lead to wasted resources for validation experiments. This risk can be mitigated by critical review of CellWhisperer-based findings and by confirming them with alternative computational approaches (such as statistical association testing) prior to costly experimentation. In contrast, we did not identify any particular risks to humans or to the environment. Given the complexity and for-research-only character of scRNA-seq profiling, it is highly unlikely that CellWhisperer results will be uncritically relied upon in clinical diagnostics and thereby harm patients. It has also been discussed whether AI tools facilitate the development of biological threats and bioweapons (Urbina et al. 2022). Given that CellWhisperer does not incorporate any biological data or knowledge that is not already in the public domain, and does not provide any dedicated functionality for the design of chemicals, viruses, or cells, we consider it highly unlikely that CellWhisperer could constitute a meaningful contribution to the toolbox of adversarial actors.

In conclusion, CellWhisperer uses AI to emulate data-centric conversations between biologists and bioinformaticians and provides an interactive tool for scRNA-seq data exploration with natural-language chats. We anticipate that natural language will become a broadly useful channel for biological data analysis and a building block for future AI-based bioinformatics research assistants that can handle diverse analytical workflows.

## Methods

### Creation of a multimodal training dataset with pairs of transcriptomes and text

To establish a large training dataset of transcriptomes and matched textual annotations, we processed two community-wide repositories: Gene Expression Omnibus (GEO) (Clough et al. 2024; Edgar et al. 2002) and CELLxGENE Census (version 2023-12-15) (CZI Single-Cell Biology Program et al. 2023).

For GEO, we obtained RNA-seq count matrices of 722,425 human transcriptomes that have been uniformly processed as part of the ARCHS^4^ project (v2.2, Date: 5-30-2023) (Lachmann et al. 2018). We removed 7,049 samples with fewer than 250 expressed genes and 9,946 samples that overlapped with our independent *disease validation dataset* (described in detail below), resulting in 705,430 transcriptomes. For each transcriptome, we obtained the associated metadata using the Entrez API with either the sample’s *experiment accession, Bio- Sample accession*, or *GEO accession* (in this order of priority), and we removed binary data as well as special characters using the *unidecode* package. We included metadata fields describing study-level descriptions such as *series design, growth protocol*, and *series_summary*, as well as sample-level fields including *sample title, treatment*, and multiple other metadata fields that varied across studies.

From CELLxGENE Census we obtained scRNA-seq count matrices of 257 studies that were conducted on human samples using one of four assays with comparable data types (10x Genomics, Seq-Well, Drop-seq, CEL-seq2). We excluded cells with fewer than 100 expressed genes, resulting in a total of 19,663,838 scRNA-seq profiles. Within each sample, we grouped cells based on cell-level metadata, using only metadata fields with fewer than 500 distinct values across all cells in the corresponding study, and we calculated pseudo-bulk (meta-cell) transcriptomes by taking the mean of the scRNA-seq count values across all cells in each metadatadefined group of cells. Each of these 376,983 pseudo-bulk transcriptomes was linked to its cell-level metadata such as the *cell type* and to study-level metadata such as a *study title* and *study abstract*.

For each sample (GEO) or pseudo-bulk transcriptome (CELLxGENE Census), we generated a concise natural-language summary from the metadata using Mixtral 8×7b (Jiang et al. 2024) (Q5_K_M quantized version), via the llama.cpp python bindings with a sampling *temperature* of 0.2, nucleus sampling *(top_p)* of 0.9, and top probability sampling *(top_k)* of 50. This LLM-based generation of concise textual annotations was guided by a prompt that we engineered based on recent best practices such as pre-action reasoning (Yao et al. 2022), roleplaying (Kong et al. 2024), and few-shot learning (Brown et al. 2020) with a manually curated set of examples. The prompt and illustrative examples of generated textual annotations are shown in **Supplementary Note 1**.

Data processing was performed on compute nodes with eight A100 80 GB GPUs. We estimate a total of 5000 GPU hours for the LLM-assisted generation of textual annotations. The training dataset is available for download via the project website (https://cellwhisperer.bocklab.org).

### Development of CellWhisperer’s multimodal embedding architecture

To enable transcriptome data analysis through natural language, we pursued a multimodal contrastive learning approach, with a neural network architecture that integrates matched pairs of transcriptomes and text into the same embedding space. Specifically, we adapted the CLIP (Contrastive Language-Image Pretraining) method (Radford et al. 2021), originally developed for joint multimodal embedding of images and text, using *pytorch* (Paszke et al. 2017) with the *lightning* (Falcon et al. 2020) and the *transformers* library (Wolf et al. 2019).

To account for the different properties of transcriptomes and natural-language text, CellWhisperer embeds the transcriptomes with Geneformer (Theodoris et al. 2023) and the metadata-derived textual annotations with BioBERT (Lee et al. 2020). The outputs of these two models are transformed into two 2048-dimensional vectors using separate adapter modules, each consisting of two learnable linear layers connected by a *ReLU* nonlinearity and followed by batch layer normalization (Shrivastava et al. 2023). To enhance computational efficiency, we adopted the LiT approach (Zhai et al. 2021), initializing both models with pre-trained weights and fine-tuning the text model as well as the adapter modules, while keeping the transcriptome model frozen.

The Geneformer model for transcriptome embedding uses 12 transformer encoder layers to process transcriptomes as “sentences of genes” ranked by their expression; it was trained on ∼30 million scRNA-seq profiles (Theodoris et al. 2023). The BioBERT model for text embedding was trained on large biomedical text corpora (Lee et al. 2020), thus providing a suitable modality-specific model for embedding the textual annotations in this study. We also tested alternative models, including scGPT (Cui et al. 2024) for transcriptome embedding and BioGPT (Luo et al. 2022) for text embedding (the CellWhisperer model and codebase allow for seamless exchange of the two modality-specific models), which led to similar results (see **Supplementary Note 2**).

### Training of CellWhisperer’s multimodal embedding model with matched pairs of transcriptomes and text

We trained our multimodal embedding of transcriptomes and text based on 1,082,413 matched pairs curated from GEO and CELLxGENE Census. For each pair, the transcriptome and textual annotation were tokenized for processing with the two modality-specific transformer models, Geneformer and BioBERT. Specifically, the transcriptomes were sorted by gene expression levels, and the top 2048 most highly expressed genes were tokenized with a dictionary of human gene symbols (Theodoris et al. 2023). The textual annotations were tokenized using WordPiece (Lee et al. 2020; Wu et al. 2016) and trimmed to a maximum of 128 tokens for training efficiency (the vast majority of textual annotations were shorter and thus remained untrimmed).

We trained the multimodal embedding model with a mini-batch size of 512 and *InfoNCE*-based loss, which maximizes the cosine similarity between matched pairs of transcriptomes and textual annotations while minimizing the cosine similarity between all other (unmatched) pairs in a given training batch. Training was scheduled for 16 epochs at a maximum learning rate of 0.00001. For the first 3% of all training steps, we froze the Geneformer and BioBERT models to only train the embedding adapters, and we linearly increased the learning rate from 0 to its maximum value (warm-up). We then unfroze the BioBERT model and continued training with a second learning rate warm-up for an additional 3% of the total number of training steps, followed by a learning rate cosine schedule over the remaining 94% of steps of the 16 epochs. The outputs of the consistently frozen Geneformer model were cached to decrease computational complexity during training.

Optimal hyperparameters, such as the maximum learning rate, were determined by stochastic grid search. As the performance metric for this optimization procedure, we tested the model’s ability to retrieve the correct textual annotation for a given transcriptome in our *disease validation dataset*. We used a deduplicated version of this dataset to increase the robustness of retrieval scoring, such as to avoid biases from data points with very similar or identical textual annotations. We also used this metric to control for overfitting during model training. The corresponding validation scores of our final model are shown in **Extended Data Fig. 1a**.

A full training run (16 epochs) was completed in less than 24 hours on a single A100 GPU. The model checkpoints are available for download on the project website (https://cellwhisperer.bocklab.org). An ablation study providing a technical evaluation of the final model is described in **Supplementary Note 2**.

### Collection and curation of validation and evaluation data

#### Validation dataset

To guide model development, we derived a thematically coherent *disease validation dataset* from GEO. We obtained 14,112 primary tissue samples that were annotated with a disease state, using a query for primary tissue and disease state to the MetaSRA database, followed by manual curation based on metadata obtained from SRA, GEO, and PubMed. To account for the effects of differences in data processing, we assessed our model on a dataset generated with a different bioinformatics pipeline and a different LLM for the validation dataset than we used for the training dataset. Specifically, we used the fetchngs and rnaseq pipelines (Ewels et al. 2020; Patel, Ewels, et al. 2024; Patel, Garcia, et al. 2024) for preparing the transcriptome data, whereas textual annotations were prepared from metadata downloaded via the Entrez API (biopython) and GPT-4 using the OpenAI API with zero-shot prompting. Because this dataset contains multiple samples with highly overlapping textual annotations, we additionally derived a *deduplicated disease validation dataset* for use in retrieval scoring. To that end, we processed all 14,112 textual annotations with BioBERT, performed hierarchical clustering on the embedding vectors (metric: cosine, linkage: average), retained the top 100 clusters, and selected for each cluster the transcriptome that was closest to the cluster center, resulting in a total of 100 deduplicated data points. We ensured the exclusion of the full validation dataset from our training data.

#### Evaluation datasets

We derived four evaluation datasets and ensured that they were fully excluded from the training data. First, we used the *Tabula Sapiens* dataset (Tabula Sapiens Consortium et al. 2022) comprising 483,152 cells across 15 donors, 24 organs, and 177 annotated cell types, with the provided *cell_ontology_class* annotation as cell type annotations (we standardized spelling/capitalization). Second, we derived a more focused *Tabula Sapiens (20 common cell types)* dataset by selecting the top 20 most common cell types in liver, lung, and blood from the full *Tabula Sapiens* dataset. Third, we created an *Immgen* dataset comprising 20 human immune cell types based on raw read counts and manually curated cell types (Heng et al. 2008; Immunological Genome Project Consortium 2023). Fourth, we included a *pancreas* dataset (dataset URL: https://figshare.com/ndownloader/files/43480497) that comprises several transcriptome profiling technologies and was previously used for benchmarking single-cell data processing (Luecken et al. 2022).

#### Demonstration dataset

To showcase CellWhisperer’s functionality for analyzing user-provided datasets, we obtained a *colonic epithelium dataset* (Parikh et al. 2019) starting from normalized read counts retrieved from GEO (GSE116222). No cell-level annotations were provided by the authors. The dataset was not included in the CELLxGENE Census and was not part of our training data.

### Evaluation of the CellWhisperer multimodal embedding model

We evaluated the trained multimodal embedding model in three complementary ways. First, we assessed CellWhisperer’s capability to predict cell properties such as cell types, tissues, and diseases in a zero-shot manner, by comparing CellWhisperer’s transcriptome embeddings with the text embeddings of the metadata-provided cell property from the evaluation datasets. To that end, we embedded the transcriptomes of each of the four evaluation datasets as well as the corresponding cell properties (as natural-language statements, for example in the following form: *‘A sample of <cell type> from a healthy individual’*). For each possible combination of the two, we quantified the agreement using the dot product (referred to as *‘CellWhisperer score’*). Finally, we *softmax*-transformed the resulting scores for each given transcriptome to obtain probabilities across property values, and we calculated the mean of ROC-AUC scores obtained for each possible property value as a metric for the model’s zero-shot cell type prediction performance.

Second, we evaluated how well the multimodal embeddings capture biological (rather than technical) differences between cells, compared to transcriptome-only embeddings. To that end, we embedded the *Tabula Sapiens* transcriptomes using either CellWhisperer’s multimodal embedding model or the transcriptome-only model (Geneformer). We then compared batch integration and cell type clustering between the two embeddings as described previously (Kedzierska et al. 2023). For cell type clustering, we used the Average Bio score (Cui et al. 2024), which is the arithmetic mean of three metrics: *average silhouette width* (Luecken et al. 2022), *normalized mutual information*, and *adjusted rank index*. For batch integration, we used a variation of the *average silhouette width* score as implemented in the *silhouette_batch* function (Luecken et al. 2022).

Third, we leveraged the broad catalog of biological phenomena covered by gene set libraries to assess which aspects of molecular biology were implicitly captured by CellWhisperer’s multimodal embedding model. To that end, we obtained 8,812 gene sets from gene set libraries of cell types (Franzén et al. 2019; Hao et al. 2021; Tabula Sapiens Consortium et al. 2022), diseases (Mckusick 1998), and Gene Ontology terms (Ashburner et al. 2000), and we performed gene set variation analysis (GSVA) using the *GSVA* package (Hänzelmann et al. 2013) with the *ssgsea* enrichment function (Barbie et al. 2009). GSVA supports gene set enrichment based on read count matrices, enabling quantitative analysis of individual samples against a background of unrelated samples. We ran GSVA across the 8,812 gene sets for the entire *disease validation dataset*, resulting in an 8,812-dimensional vector of gene set enrichments for each transcriptome. For the same transcriptomes, we also derived CellWhisperer scores by embedding the textual annotations of the gene sets (e.g. “colorectal cancer” from the *OMIM_Extended* disease library or “Response To Type I Interferon” corresponding to *GO:0071357 ID* from *GO_Biological_Process_2023*). We then compared the GSVA and CellWhisperer scores for each gene set across the transcriptomes in the dataset using the Pearson correlation and Kolmogorov- Smirnoff statistic (included here for its robustness to non-linear CellWhisperer scores).

### Development and training of CellWhisperer’s multimodal LLM for natural-language chats

To enable natural-language chats with CellWhisperer’s multimodal embedding model, we fine-tuned the Mistral 7B LLM based on data-centric conversations about individual transcriptomes and cell states. We generated 106,610 such conversations for transcriptomes from our training dataset of 1,082,413 GEO and CELLxGENE Census data points. To mitigate historical bias in these community repositories, we took a weighted subsample, inversely proportional to the local point density calculated with *densMAP* (Narayan et al. 2021), such that transcriptome-text pairs in lowly covered regions were preferentially picked. For each transcriptome, we generated a chat-like conversation using one of two LLMs (GPT-4 or Mixtral) on the basis of the transcriptome’s top 50 most highly expressed genes, the top 50 GSVA-derived gene sets, and the transcriptome’s textual annotation. We prepared conversations in four ways, resulting in *simple, detailed, complex*, and *conversational chats*. The LLM prompts and examples of the generated conversations are shown in **Supplementary Note 1**.

- *Simple chats* were generated for 10,000 transcriptomes by using a generic question (randomly selected from a list of 10 manually prepared candidates: “What does the sample represent?”, “What do these cells represent?”, etc.) and the transcriptome’s generated textual annotation as the answer.
- *Detailed chats* were generated for 10,000 transcriptomes using an LLM (GPT-4 via the OpenAI API) with zero-shot prompting and the same questions as in the *simple chats*. GPT-4 produced much more extensive answers as compared to the transcriptome’s textual annotations used in the *simple chats*.
- *Complex chats* were generated for 5,000 transcriptomes using an LLM (GPT-4 via the OpenAI API) with a few-shot prompt using pre-action reasoning to produce a single profound question-answer pair.
- *Conversational chats* for 81,610 transcriptomes were generated using an LLM (Mixtral 8×7b) with a few-shot prompt that instructed a natural conversation with multiple questions and answers between a researcher and an AI assistant about the biological state of the assayed sample or cell cluster.

Based on these training conversations, we implemented CellWhisperer’s conversational capabilities following the LLaVA approach (H. Liu et al. 2023). We initialized a two-layer adapter module that transforms the 2048dimensional *CellWhisperer* transcriptome embedding into eight 4096-dimensional embeddings, corresponding to eight token embeddings within the Mistral 7B LLM architecture (Jiang et al. 2023), which is the basis for CellWhisperer’s multimodal LLM. To train our model based on the training conversations, we passed transcriptome-derived token embeddings alongside their corresponding questions to the LLM and optimized the LLM and the transcriptome-token adapter to generate the corresponding answers using the original auto-regressive learning objective of the Mistral LLM, with a loss mask for the question parts of the conversations. In a first training step, we kept the LLM frozen and trained the adapter layers with supervision on an extended version of the *simple chat* conversations that included our entire training dataset (1,082,213 question-answer pairs) for one epoch. In a second training step, we unfroze the Mistral LLM and fine-tuned both the LLM and the adapter layers on the 106,610 generated training conversations for one epoch.

LLM fine-tuning was performed on four A100 80 GB GPUs with a total runtime of three hours. The generated conversation datasets and model checkpoints are available for download on the project website.

### Evaluation of CellWhisperer’s multimodal LLM for natural-language chats

To establish an evaluation dataset for CellWhisperer’s natural-language chat functionality, we randomly selected 200 of the 106,610 training conversations, half derived from GEO (bulk transcriptomes) and the other half from CELLxGENE Census (pseudo-bulk transcriptomes based on scRNA-seq data) and with proportional representation of *simple, complex*, and *conversational chats*. This *conversation evaluation dataset* was excluded from all training and fine-tuning to allow independent evaluation. The conversations were trimmed to the first question-answer pair, and passages that did not refer to biological cell states were manually removed.

We first evaluated whether, for a given conversation, CellWhisperer’s multimodal LLM preferred the correctly matched transcriptome over randomly mismatched transcriptomes. We quantified this preference using the perplexity metric, which is defined as the exponentiated average negative log-likelihood of the sequence of tokens that correspond to the “ground truth” answer for a given prompt (i.e., a question and a matched or unmatched transcriptome embedding). In other words, perplexity measures the degree of surprise for the model when confronted with an answer to a given pair of a question and a matched or unmatched transcriptome.

To contextualize these perplexity measurements, which are both model-specific and task-specific and not readily interpretable as absolute numbers, we computed the perplexity for each test conversation not only with the matched transcriptome embedding but also with 30 randomly sampled transcriptome embeddings. Against this background, we computed the perplexity quantile of the correctly matching transcriptome embedding as a metric of how well CellWhisperer took the transcriptome data into account when generating its chat response.

As a separate line of validation, we assessed whether CellWhisperer’s multimodal LLM could correctly predict cell types in the *Tabula Sapiens (20 common cell types)* dataset. For each of the 20 cell types, we sampled 20 pseudo-bulk transcriptomes, resulting in a total of 400 individual transcriptomes. We asked a simple question (*‘Which cell type is this cell?’*) and converted the annotated cell type label to a natural-language answer by prefixing them with the following text: *‘This cell is a ‘*. To evaluate the preference of a given transcriptome for its annotated cell type, we calculated the perplexity for all possible cell-type answer texts and determined the quantile of the correct cell-type perplexity against the background of all unmatched-answer perplexities.

### Integration of CellWhisperer into CELLxGENE Explorer, web hosting, and user-provided dataset processing

To make CellWhisperer easily accessible and broadly useful, we integrated it into the popular CELLxGENE Explorer software (v1.2.0) for interactive analysis of scRNA-seq data (Megill et al. 2021). An introduction to the CELLxGENE Explorer is available on the project website (https://docs.cellxgene.cziscience.com). We extended the CELLxGENE Explorer user interface with a chat box and implemented two core API endpoints: (i) natural-language chat functionality to get information about a selected group of cells; (ii) a search interface to obtain *CellWhisperer* scores for user-provided free-text queries, which are displayed as cell-level color maps. We flexibly scan and route the free-text user requests to these two endpoints with a regular expression that matches the first word to be one of “show”, “show me”, “search” or “search for”. The CellWhisperer chats inside CELLxGENE Explorer are primed with a hidden conversation initiation that provides the 50 top genes for the selected cells/samples (normalized expression across all transcriptomes in the dataset, to improve the representation of lowly expressed but highly relevant genes such as transcription factors).

We host the CellWhisperer-augmented version of CELLxGENE Explorer as a web application using *docker / docker-compose*. Each dataset receives its dedicated server job, which is jointly exposed via an *nginx* web server. CellWhisperer’s multimodal embedding and LLM models are hosted via independent API server jobs, which are accessed by all dataset-specific servers. This way, the model is loaded only once into GPU RAM, and it allows running the CellWhisperer software locally on computers without a GPU. User-provided datasets are preprocessed for interactive analysis using an automated pipeline. This includes the computation of CellWhisperer transcriptome embeddings and 50 top genes for each cell in the dataset. Next, a UMAP is calculated on the transcriptome embeddings, followed by clustering using the Leiden algorithm and the LLM-based generation of a brief textual annotation for each cluster. To that end, we use CellWhisperer’s multimodal LLM on the cluster-averaged transcriptome embedding to generate detailed textual descriptions. These texts are then condensed into brief textual annotations using an LLM such as GPT-4 (via the OpenAI API, used in this study) or Mixtral 8×7B (which can be run locally, supported by the source code). All prompts are provided in **Supplementary Note 1**. The preprocessed user-provided dataset is stored as a single h5ad object that is readily loaded into CellWhisperer. We provide this data processing pipeline as a single shell command and used it to process the datasets in **Fig. 1, Fig. 3**, and **Fig. 4**, and for our demonstration video (https://cellwhisperer.bocklab.org). Data processing has a runtime in the order of minutes for typical datasets on a compute node with a single A100 GPU, or in the order of hours when relying on a conventional quad-core CPU-only computer.

## Data availability

CellWhisperer was trained on publicly available datasets from multiple sources, including GEO and CELLxGENE Census. These datasets are available from their original sources as documented in the Methods section and the CellWhisperer source code, and our software pipelines support the automatic download of the required data. Moreover, the LLM-curated textual annotations that were used for training and model evaluation are available for download from the project website (https://cellwhisperer.bocklab.org).

## Code availability

The source code underlying this project is available on GitHub (https://github.com/epigen/cellwhisperer). This includes the CellWhisperer codebase and a version of CELLxGENE Explorer with CellWhisperer integration. We further provide automatic start-to-finish pipelines to reproduce all analyses in this study and to process user-supplied datasets for interactive analysis using a local copy of the CELLxGENE Explorer with CellWhisperer integration. Where we build on external codebases (Geneformer, LLaVA, CELLxGENE Explorer), we forked, adapted, and integrated them as submodules in the CellWhisperer GitHub repository. In addition to the CellWhisperer source code, we provide the trained CellWhisperer model as a PyTorch Lightning checkpoint for download on the project website (https://cellwhisperer.bocklab.org). Detailed instructions on installing CellWhisperer and reproducing this study are provided in the GitHub repository.

## Author Contributions (following the CRediT taxonomy)

Conceptualization, M.S., C.B., P.P.; Methodology, M.S., P.P., M.P., A.H., V.S., D.M.; Software, M.S., P.P., D.M., J.B.; Validation, P.P., M.S., M.P., T.K.; Resources, C.B., J.M., E.T.; Data Curation, M.S., D.M., P.P., M.P.; Writing – Original Draft, M.S., C.B., P.P.; Writing – Review & Editing, all authors; Visualization, M.S., P.P., D.M.; Supervision, C.B., E.T., J.M.; Project Administration, M.S.; Funding Acquisition, C.B., E.T., J.M.

## Acknowledgments

We thank the members of the Bock lab and the AI Institute of the Medical University of Vienna for their help and advice. CB is supported by a European Research Council (ERC) Consolidator Grant (no. 101001971). EMT is supported by the Austrian Science Fund (P34958) and by an ERC Consolidator Grant (no. 101087883). CB and EMT are supported by the Vienna Science and Technology Fund (WWTF) (LS18-049, LS20-045).

## Extended Data Figures

**Extended Data Figure 1.**
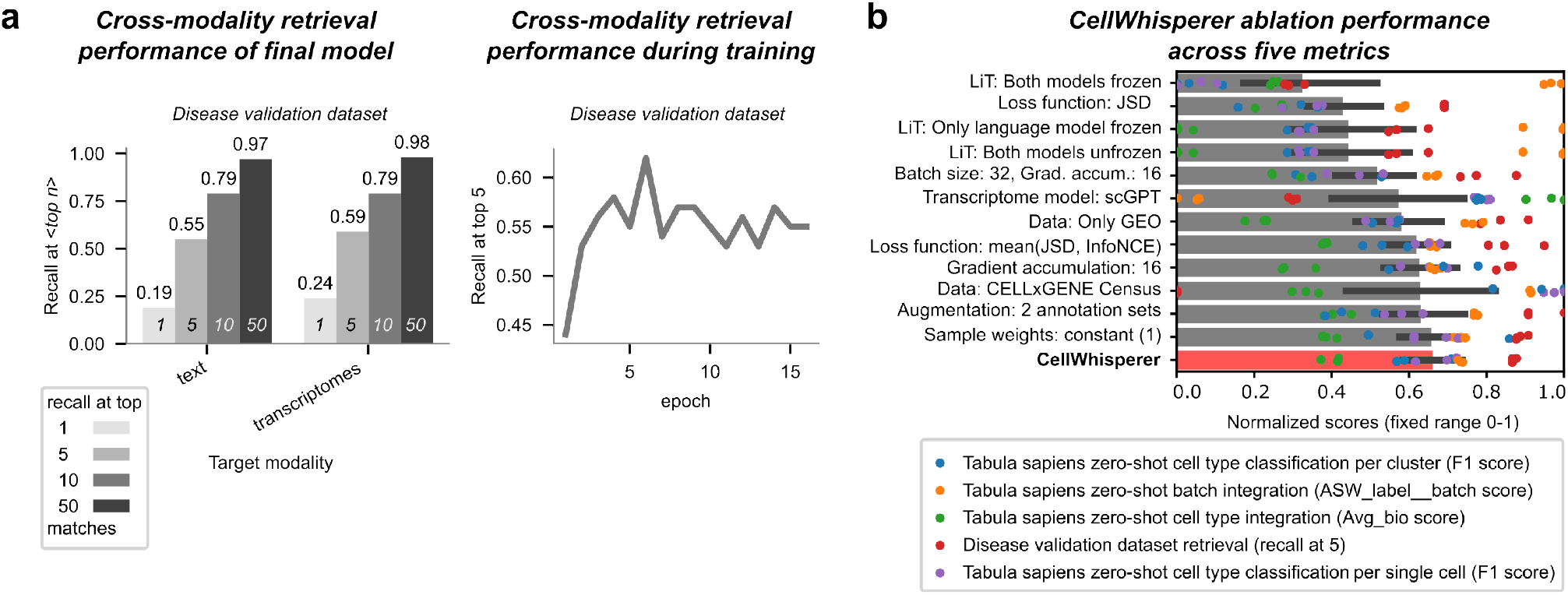
CellWhisperer cross-modality retrieval performance and model ablation testing. a) Cross-modality retrieval performance of CellWhisperer’s multimodal embedding model, evaluated on the deduplicated disease validation dataset for the trained model (left) and during training (right). b) Performance comparison across alternative architecture and hyperparameter choices in an ablation study of the CellWhisperer embedding model. For each alternative model, the mean performance across five metrics is shown, each based on three models trained with different seeds. The values for each metric (shown as dots) were min-max normalized across models. F1 scores are macro averages across labels. The bars indicate the mean across all data points, with error bars corresponding to 95% confidence intervals. The final CellWhisperer embedding model (marked in bold) was trained with batch size 512, no gradient accumulation, frozen Geneformer transcriptome model, InfoNCE loss, GEO and CELLxGENE Census datasets, density-based sample weighting, and no data augmentation. JSD: Jensen-Shannon divergence information maximization loss. InfoNCE: information noise contrastive estimation.

**Extended Data Figure 2.**
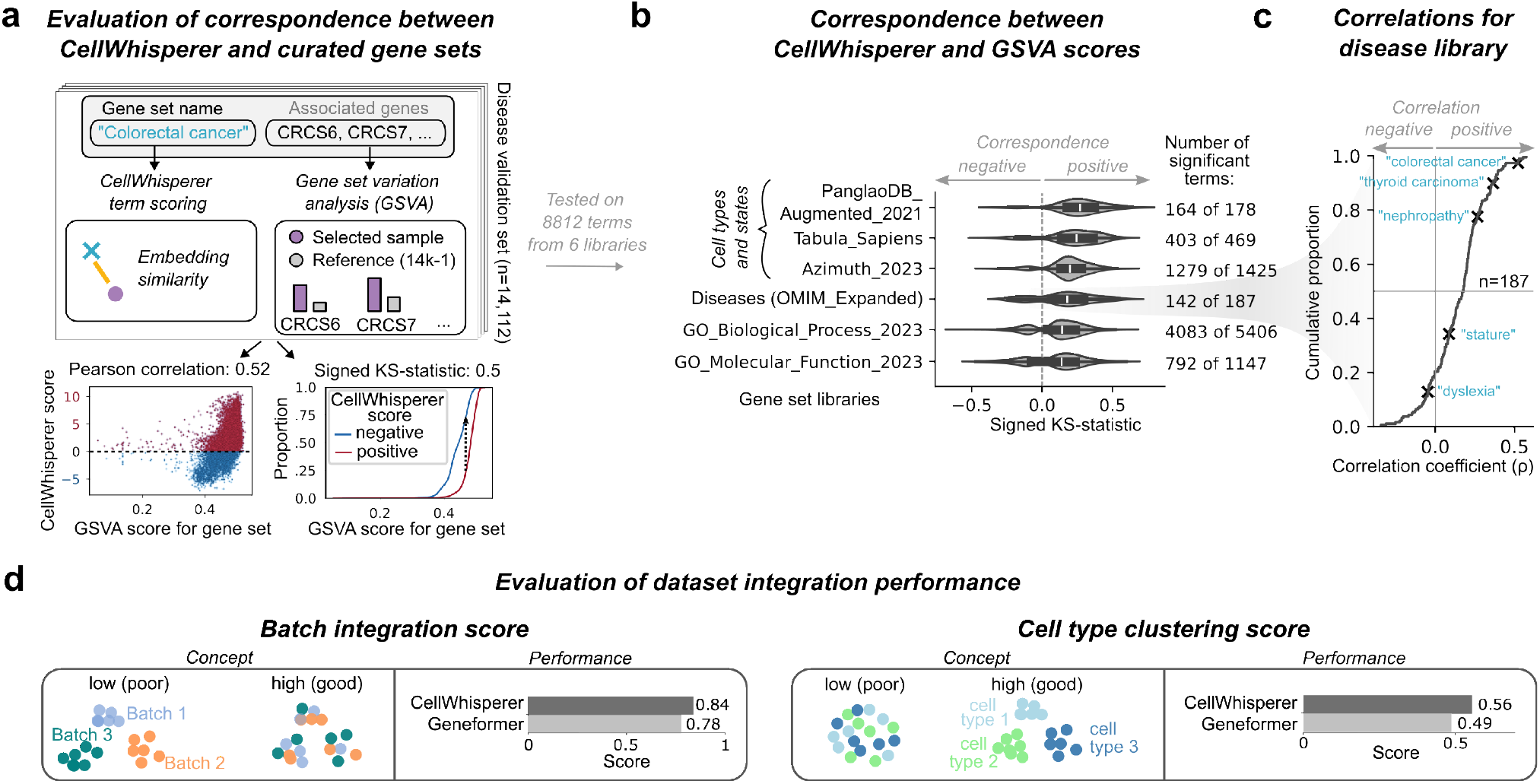
CellWhisperer batch integration performance and evaluation based on gene sets. a) Benchmarking of CellWhisperer scores using gene set labels as textual annotations of biological functions versus the enrichment of the associated genes in the transcriptome dataset (GSVA scores). The plots at the bottom show the Pearson correlation and Kolmogorov-Smirnoff (KS) statistic for the CellWhisperer and GSVA scores across the 14,112 samples of the disease validation dataset, for the example gene set “colorectal cancer”. b) Correspondence (as measured by the signed KS statistic) between CellWhisperer and GSVA scores broken down by gene set library. The number of gene sets with significant distribution differences in the expected direction is indicated on the right (significance threshold: 0.05). Standard violin plots are shown with inner boxplots corresponding to the interquartile range and whiskers extending to the farthest data point within 1.5 times the interquartile range. c) Distribution of Pearson correlation coefficients between CellWhisperer and GSVA scores for the disease gene set library OMIM_Expanded. Selected terms are labeled for illustration. d) Illustration of the batch integration and cell type clustering tasks for performance comparison between CellWhisperer’s multimodal model and its underlying transcriptome-only Geneformer model. Transcriptomes from multiple batches and donors within the Tabula Sapiens dataset were embedded and subsequently examined on how well the technical variation was removed (batch integration score) and how well the biological differences between cell types were retained (cell type clustering score).

**Extended Data Figure 3.**
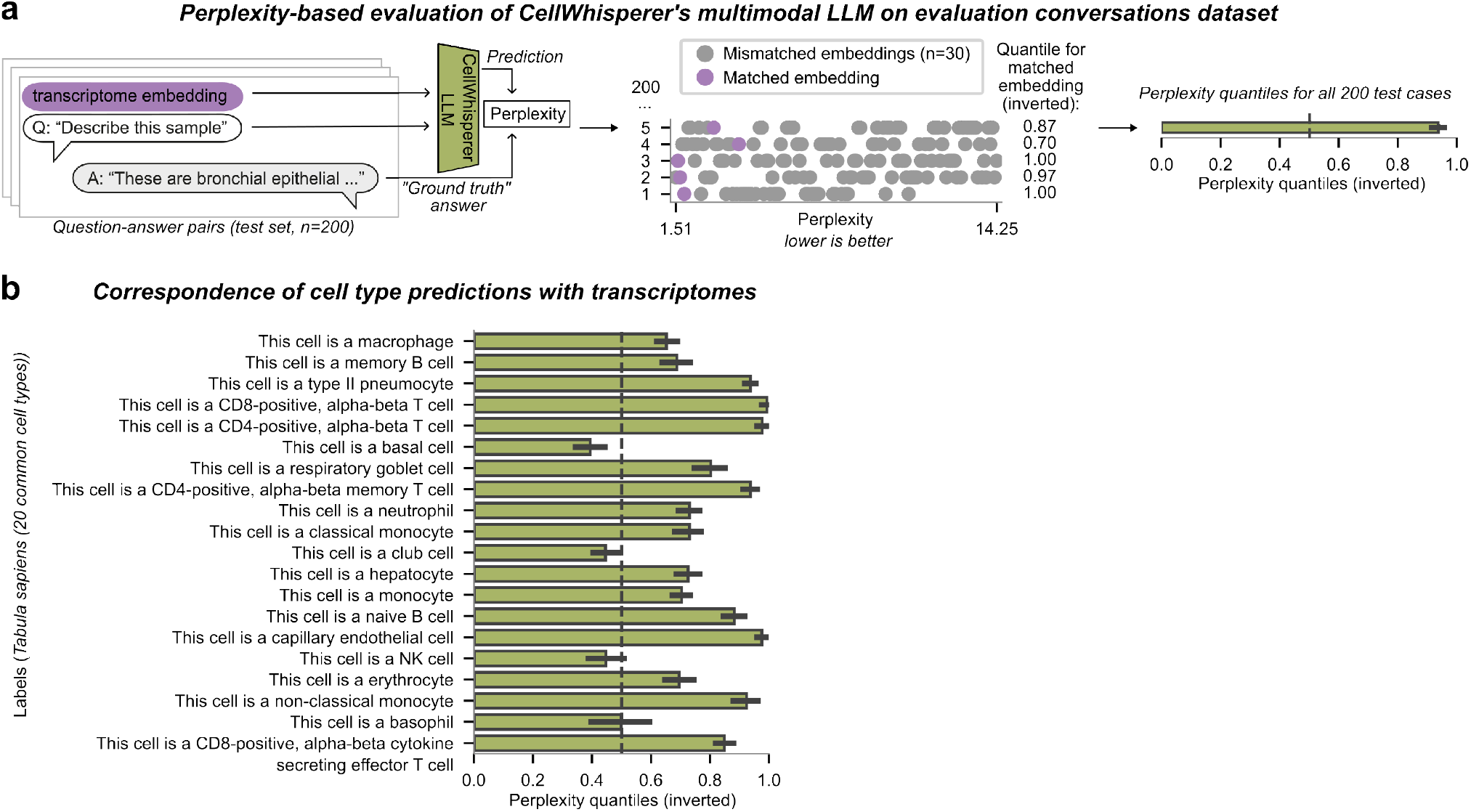
Evaluation of CellWhisperer text generation and chat performance. a) Perplexity-based evaluation of CellWhisperer’s multimodal LLM across 200 question-answer pairs from the conversation evaluation dataset with matched versus mismatched transcriptome embeddings. For each question-answer pair, the plot shows the inverted quantile of the perplexity for the matched transcriptome embedding relative to the mean perplexity across 30 mismatched embeddings. Higher values indicate better performance, and the dashed line at 0.5 indicates the random baseline. b) Perplexity-based performance evaluation of cell type prediction by the CellWhisperer multimodal model for the Tabula Sapiens (well-annotated cell types) dataset. Quantiles were computed by comparing the perplexity of the correct cell type response against 30 mismatched responses for each transcriptome. Bars indicate the mean across all data points, and error bars correspond to the 95% confidence interval.

## Supplementary Information

*Supplementary Note 1*: Prompts and examples for LLM-assisted curation of the CellWhisperer training data

*Supplementary Note 2*: Ablation study of the CellWhisperer multimodal model

*Supplementary Table 1:* Statistical evaluation of gene set correspondence between CellWhisperer and GSVA

