## Supplementary Note 1 for "Multimodal learning of transcriptomes and text enables interactive single-cell RNA-seq data exploration with natural-language chats"

### Supplementary Note 1: Prompts and examples for LLM-assisted curation of the CellWhisperer training data

Training of the CellWhisperer models relies on consistently processed transcriptome metadata for the generation of concise natural-language annotations and chat conversations using large language models (LLMs). In this Supplementary Note, we list all LLM prompts that were used for training data curation, together with examples of input data and the corresponding LLM-curated outputs. For each of the LLM data curation tasks we provide three types of information: (i) The prompt; (ii) example input data (for consistency, we rely on the same input sample across the different processing steps, which is listed below only once for all prompts); (iii) the LLM-generated output. Further methodological details are provided in the **Methods** section of the paper.

#### 1. Principles of prompt engineering for CellWhisperer training data curation

LLMs are known to depend on well-optimized prompts to achieve best results. We followed an iterative prompt engineering approach, the results of which are shown below, relying on several recently established techniques. First, as commonly observed in deployed LLMs such as ChatGPT, responses benefit if the LLM first reiterates the question in an ordered form and recapitulates established facts about the topic; this has been referred to as pre-action reasoning (Yao et al. 2022). We employ this technique to obtain concise final outputs, as we instruct the LLM to first gather all relevant information and then aggregate it into a compressed response. Second, it is often helpful to assign a hypothetical role to the LLM, encouraging it to respond in the voice of an expert for the task at hand (e.g., "You are a biomedical researcher"); this has been shown to improve the quality of responses (Kong et al. 2024). Third, to illustrate and further contextualize the instructions in our prompt, we provide the LLM with hand-curated examples of how we expect it to transform inputs into outputs, by appending multiple such pairs to the prompt, a technique referred to as few-shot learning (Brown et al. 2020).

#### 2. Generation of textual annotations from sample metadata

*Prompt for deriving concise textual annotations from sample metadata (LLM: Mixtral 8x7B with nine few-shot learning examples, which are available in source code repository)*

```
You are a biomedical researcher. Your task is to extract the biologically relevant information
from a sample's metadata indicated by <<<>>> and representing them in natural language.

# Instructions

## Represent Biological Information
Only consider information that are reflective of the sample's biology, i.e. the phenotype and
transcriptome of the contained cells, such as cell type or any pertinent conditions or
treatments.

## Study-Level Information
```

```
This is just additional context and should not be included in the response, beyond informing
about the the nature of the sample.

## Ignore Non-Biological Information
Ignore information that is not relevant to the cellular state of the sample. Specifically,
exclude all IDs and details about technical and methodological processes such as sequencing type,
read length, platform, or any other laboratory technique-related information.

## Response Template
Respond in two parts following the JSON template below.
{{
    "1. Relevant Information to be Included": <List[str]: List of information to be extracted,
according to the instructions above>,
    "2. Final Response": <str: brief natural language descripton of the sample focusing solely on
the biological aspects that contribute to understanding its cellular state>,
}}

# Examples
Here are some examples:

{few_shot_block}

# Provided Metadata

<<<
{query}
>>>
```

*Input example of metadata for a bulk transcriptome sample obtained from GEO/SRA with ID SRX7552910*

- geo_title: T2-4
- geo_source_name: Renal Cell Carcinoma
- raw_biosample_metadata: source_name: Renal Cell Carcinoma; tissue: Renal Cell Carcinoma; tissue preparation: nephrectomy specimens preserved in FFPE; metastatic: No; tumor stage: 2; sex: male
- treatment:
- treatment_protocol: None
- series_summary: Presented is the differential expression of mRNA and miRNA in 44 Stage I and Stage II patients who developed metastasis within 5 years of nephrectomycompared to 21 patients who remained disease free. Extracted RNA from nephrectomy specimens preserved in FFPE blocks was sequenced using RNAseq. MiRNA expression was done using the TaqMan OpenArray qPCR protocol.
- series_design: Examination of molecular differences between metatstatic versus non-metatstatic in stage I and stage II
- growth_protocol: None

*Textual annotation output generated based on the metadata for bulk RNA-seq sample with ID SRX7552910*

Renal cell carcinoma tissue sample taken from a male patient at stage 2, with no metastasis, preserved in FFPE blocks

*Input example of metadata for scRNA-seq dataset obtained from CELLxGENE Census with ID 1e6a6ef9-7ec9-4c90-bbfb-2ad3c3165fd1) and meta-cell number 3725*

- cell_type: club cell
- development_stage: unknown
- organism: Homo sapiens
- self_reported_ethnicity: unknown
- sex: male
- suspension_type: cell
- tissue: lung

- study_description: Non-small cell lung cancer (NSCLC) is characterized by molecular heterogeneity with diverse immune cell infiltration patterns which has been linked to therapy sensitivity and resistance. However, full understanding of how immune cell phenotypes vary across different patient subgroups is lacking. Here, we dissect the NSCLC tumor microenvironment at high resolution by integrating 1,283,972 single cells from 556 samples and 318 patients across 29 datasets, including our dataset capturing cells with low mRNA content. We stratify patients into immune-deserted, B cell, T-cell, and myeloid cell subtypes. Using bulk samples with genomic and clinical information, we identify cellular components associated with tumor histology and genotypes. We then focus on the analysis of tissue-resident neutrophils (TRNs) and uncover distinct subpopulations that acquire new functional properties in the tissue microenvironment, providing evidence for the plasticity of TRNs. Finally, we show that a TRN-derived gene signature is associated with anti-programmed cell death ligand 1 (PD-L1) treatment failure.,
- study_title: The single-cell lung cancer atlas (LuCA) -- extended atlas
- sample_source_name: Chen_Zhang_2020_NSCLC-5
- ann_coarse: Epithelial cell
- ann_fine: Club
- cell_type_major: Club
- cell_type_neutro: Club
- cell_type_neutro_coarse: Club
- cell_type_predicted: Epithelial cell malignant
- cell_type_tumor: Club
- dataset: Chen_Zhang_2020
- doublet_status: singlet
- origin: tumor_primary
- origin_fine: tumor_primary
- platform: 10x
- study: Chen_Zhang_2020
- tumor_stage: advanced
- uicc_stage: III

*Textual annotation output generated based on the metadata for scRNA-seq sample with ID 1e6a6ef9-7ec9-4c90-bbfb-2ad3c3165fd1 and meta-cell number 3725*

Club cells derived from a primary lung tumor of a male individual with advanced stage non-small cell lung cancer (NSCLC), UICC stage III.

### 3. Generation of question-answer conversations for CellWhisperer's multimodal LLM

*Hand-crafted questions for the "simple" and "detailed" conversations*

- *What does the sample represent?*
- *What does the transcriptome represent?*
- *What do these cells represent?*
- *Give a brief description of the sample.*
- *Give a brief description of these cells.*
- *Present a compact description of the sample's key features.*
- *Summarize the state of these cells.*
- *Provide a brief description of these cells.*
- *Provide a brief description of the given sample.*
- *Describe the sample concisely.*
- *Describe these cells concisely.*

*Prompt for "complex" conversations (GPT-4 with three few shot examples provided in source code repository)*

```
You are an AI molecular analyst with a deep understanding of cells and their states. Your task is
to generate a question and a response that delve into the intricacies of a biological sample of
human cells.

# Instructions

## Provided information
You are equipped with a detailed description of a biological sample of cells, including
- a succinct natural language summary of the biological state of the sample
- a prioritized list of the most prominently expressed genes (with the highest expression levels
listed first)
- a prioritized list of the most active pathways, processes and other ontological terms (with the
most significant ones listed first)

## First contemplate
First contemplate about the provided information to obtain a biological intuition of the nature o
the provided sample. Use this intuition (rather than the provided raw annotation and gene lists)
to identify an interesting biological question with a matching insightful, but concise, response.
Finish the contemplation with a suggestion for the topic of the question.

## The researcher's question
The researcher possesses knowledge in the biomedical domain (such as molecular biology,
immunology, and cancer biology), but is unfamiliar with the specifics of the provided cellular
sample. They are interested in the sample's phenotype, cell type/state, origin,
treatment/perturbation, active processes/pathways and pose a specific and interesting question
that will unveil any of the contemplated information. Examples for interesting questions:
- How have these cells been treated?
- What are the biological processes and pathways that stand out in this sample?
- Does this sample exhibit pathogenic characteristics?

## The AI system's response
The AI system provides a comprehensive, concise and insightful response to the researcher's
question, drawing from summary in the 'contemplation' step, rather than from the raw provided
information. As the researcher does not have any information about the sample, the response needs
to be phrased as such. Specifically, the AI response should sound like it is directly
investigating the biological state of the cellular sample, without having any additional
information of its origin or processing steps. In line with this, all provided information should
explicitly acknowledge the uncertainty of the AI system (e.g. use phrases like \"it seems\",
\"could be\", \"likely\", \"potentially\", etc.).

## Format
Generate the dialogue as a JSON array, following this schema:

[{\"contemplation\": \"<str: contemplation of the sample's state>\"}, {\"from\": \"researcher\",
\"value\": <str: the researcher's question>}, {\"from\": \"ai\", \"value\": <str: the AI system's
response>}]
```

*Input example for "complex", "detailed", and "conversation" conversations (based on data for SRX7552910)*

```
Sample annotation: Renal cell carcinoma tissue sample taken from a male patient at stage 2, with
no metastasis, preserved in FFPE blocks.

Gene sets (e.g. pathways, GO terms): Bone Marrow-hematopoietic Stem Cell, Blood-hematopoietic Stem
Cell, Blood-cd141-positive Myeloid Dendritic Cell, Bone Marrow-myeloid Progenitor, Blood-basophil,
Mammary-pericyte Cell, Mammary-vein Endothelial Cell, Thymus-innate Lymphoid Cell, Skin-nkt Cell,
Skin-cd4-positive, Alpha-Beta Memory T Cell, Skin-naive Thymus-Derived Cd4-Positive, Alpha-Beta T
Cell, Skin-naive B Cell, Blood-myeloid Progenitor, Spleen-cd1c-positive Myeloid Dendritic Cell,
Blood-plasmacytoid Dendritic Cell, Spleen-naive Thymus-Derived Cd4-Positive, Alpha-Beta T Cell,
Skin-t Cell, Large Intestine-cd4-positive, Alpha-Beta T Cell, Skin-cd8-positive, Alpha-Beta Memory
T Cell, Skin-cd4-positive Helper T Cell, Skin-nk Cell, Lung-smooth Muscle Cell, Mammary-endothelial
Cell Of Artery, Large Intestine-b Cell, Blood-naive Thymus-Derived Cd4-Positive, Alpha-Beta T Cell,
Lung-cd4-positive, Alpha-Beta T Cell, Blood-plasma Cell, Mammary-b Cell, Bone Marrow-cd4-positive,
```

```
Alpha-Beta T Cell, Small Intestine-cd4-positive, Alpha-Beta T Cell, Pancreas-b Cell, Salivary Gland-cd4-positive Helper T Cell, Thymus-dn3 Thymocyte, Eye-t Cell, Mammary-t Cell, Bone Marrow-memory B Cell, Thymus-memory B Cell, Lymph Node-regulatory T Cell, Lung-bronchial Vessel Endothelial Cell, Fat-leucocyte, Small Intestine-cd8-positive, Alpha-Beta T Cell, Vasculature-b Cell, Skin-cd8-positive, Alpha-Beta Cytotoxic T Cell, Lymph Node-innate Lymphoid Cell, Trachea-cd4-positive, Alpha-Beta T Cell, Fat-t Cell, Thymus-vein Endothelial Cell, Spleen-cd4-positive, Alpha-Beta Memory T Cell, Thymus-dn1 Thymic Pro-T Cell, Large Intestine-mast Cell

Top expressed normalized genes: NEB, MUC12, C4A, IGHG1, IGHG1, SCARNA10, SNORD3D, IGLC1, IGFN1, CTAGE9, IGHG2, UGT2B7, SCARNA2, IGLC2, VWF, IGHGP, ENSG00000243302, USP17L15, NAIPP4, FLG, COL3A1, IGLC3, SCARNA7, PKHD1, ENSG00000284454, HRNR, LRRC37A, ARHGEF34P, COL1A2, ENSG00000281380, NAIPP2, ARHGEF35, IGKC, MGAM, MGAM, RGPD1, AHNAK2, NPIPB12, PREX2, PPIAP22, LRRC37A3, ENSG00000215156, SST, SCARNA5, SCARNA5, DNAH11, RGPD3, ENSG00000283738, USP17L6P, C3
```

*Generated output for the "complex" conversation prompt and the question-answer input for SRX7552910*

- Contemplation: This sample is from a stage 2 renal cell carcinoma (RCC) tissue of a male patient, preserved in formalin-fixed paraffin-embedded (FFPE) blocks. The absence of metastasis suggests localized disease. The gene expression and pathway enrichment reflect a complex interaction between tumor cells and the tumor microenvironment, including immune infiltration and extracellular matrix remodeling. High expression of genes such as VWF (involved in blood clotting and endothelial cell function), COL3A1, COL1A2, and COL1A1 (collagen genes involved in extracellular matrix structure), alongside MUC12 (a mucin gene potentially indicative of a protective barrier or altered cell signaling) suggests active angiogenesis and tissue remodeling. The presence of immune-related pathways and cells indicates an active immune response or immune evasion mechanisms. The gene expression profile points towards a tumor microenvironment that is actively engaged in maintaining the tumor structure and potentially facilitating its growth, despite the localized nature of the disease. The specific characteristics of this RCC sample, such as its stage, cell interactions, and potential vulnerabilities, could be the focus of an interesting question.
- Question (from researcher): What does the gene expression profile indicate about the tumor microenvironment in this RCC sample?
- Response (from AI assistant): The gene expression profile of this stage 2 renal cell carcinoma sample suggests a dynamic and complex tumor microenvironment. There's evidence of significant extracellular matrix remodeling, indicated by the high expression of collagen genes like COL3A1, COL1A2, and COL1A1, which are essential for tissue structure and integrity. The presence of VWF hints at active angiogenesis, which is crucial for tumor nourishment and growth. Additionally, the expression of MUC12 may suggest altered cell signaling or a protective barrier function in the tumor cells. Immune-related pathways and gene sets suggest either an active immune response within the tumor microenvironment or mechanisms of immune evasion by the tumor. Together, these factors underscore the interactions between tumor cells, the extracellular matrix, and the immune system, contributing to the tumor's growth and potential resistance to therapy.

*Prompt for "detailed" conversations (LLM: GPT-4, zero-shot)*

```
You are an AI molecular analyst with a deep understanding of cells and their states. Your task is
to generate a question and a response that delve into the intricacies of a biological sample of
human cells.

# Instructions

## Provided information
You are equipped with a detailed description of a biological sample of cells, including
- a succinct natural language summary of the biological state of the sample
- a prioritized list of the most prominently expressed genes (with the highest expression levels
listed first)
- a prioritized list of the most active pathways, processes and other ontological terms (with the
most significant ones listed first)
```

```
## First contemplate
First contemplate about the provided information to obtain a biological intuition of the nature o
the provided sample. Use this intuition (rather than the provided raw annotation and gene lists)
to identify an interesting biological question with a matching insightful, but concise, response.
Finish the contemplation with a suggestion for the topic of the question.

## The researcher's question
The researcher possesses knowledge in the biomedical domain (such as molecular biology,
immunology, and cancer biology), but is unfamiliar with the specifics of the provided cellular
sample. They are interested in the sample's phenotype, cell type/state, origin,
treatment/perturbation, active processes/pathways and poses the following question: {question}

## The AI system's response
The AI system provides a comprehensive, concise and insightful response to the researcher's
question, drawing from summary in the 'contemplation' step, rather than from the raw provided
information. As the researcher does not have any information about the sample, the response needs
to be phrased as such. Specifically, the AI response should sound like it is directly
investigating the biological state of the cellular sample, without having any additional
information of its origin or processing steps. In line with this, all provided information should
explicitly acknowledge the uncertainty of the AI system (e.g. use phrases like \"it seems\",
\"could be\", \"likely\", \"potentially\", etc.).

## Format
Generate the dialogue as a JSON array, following this schema:

[{{\"contemplation\": \"<str: contemplation of the sample's state>\"}}, {{\"from\":
\"researcher\", \"value\": <str: the researcher's question>}}, {{\"from\": \"ai\", \"value\":
<str: the AI system's response>}}]
```

*Generated output for the "detailed" conversation prompt and the question-answer input for SRX7552910*

- Contemplation: The provided sample is from renal cell carcinoma tissue, indicating a cancerous state. The gene expression profile and active pathways suggest a complex interplay of various cell types, including hematopoietic stem cells, myeloid and dendritic cells, T cells, B cells, and endothelial cells, among others. This diversity implies a rich microenvironment within the tumor, potentially reflecting immune infiltration, angiogenesis, and possibly the presence of cancer stem cells. The high expression of genes like NEB, MUC12, and C4A, among others, could be indicative of specific cancer-related processes such as cell adhesion, immune response modulation, and extracellular matrix remodeling. The presence of genes associated with hematopoiesis and immune cells suggests an active immune response or infiltration within the tumor microenvironment. The challenge lies in deciphering the roles these diverse cell types and pathways play in tumor progression, immune evasion, and their potential as therapeutic targets or biomarkers.
- Question (from researcher): Describe these cells concisely.
- Response (from AI assistant): The sample appears to be from a renal cell carcinoma at stage 2, characterized by a complex tumor microenvironment. This environment includes a variety of cell types such as hematopoietic stem cells, myeloid and dendritic cells, T cells, B cells, and endothelial cells, suggesting immune infiltration and angiogenesis. The gene expression profile, with genes like NEB, MUC12, and C4A highly expressed, points towards processes such as cell adhesion, immune response modulation, and extracellular matrix remodeling. These features could reflect the tumor's interaction with its microenvironment, including immune evasion mechanisms and potential angiogenesis. The diversity of cell types and active pathways indicates a rich, dynamic interplay within the tumor, which could have implications for therapy and prognosis.

*Prompt for "conversation" conversations (Mixtral, five few shot examples provided in source code repository)*

```
You are a molecular analyst and play writer with a deep understanding of cellular states and
phenotypes. Your task is to generate a realistic conversation between a biomedical researcher and
an AI assistant, chatting about a biological sample of human cells.

# Instructions
```

```
## Provided information
You are given a description of a biological sample of cells, that represents its state/phenotype.
This description includes
- a concise natural language annotation of the biological state of the sample
- a ranked list of the most strongly expressed genes (most upregulated genes come first)
- a ranked list of the most active pathways and enriched gene ontological terms (most signifcant
ones come first)

## The researcher's questions
The researcher has an interest in molecular biology, immunology, cancer biology, etc. and
initially doesn't know anything about the provided biological sample. He asks questions to learn
about the biological nature of the sample. The questions should be relevant, diverse, addressable
(given the provided information) and in the natural tone of a scientist curious about the
nature/biology of the sample. Specifically the researcher uses simple and conversational
language, asking questions on the biology of the sample (e.g. active pathways, processes,
mechanisms, regulation patterns etc.). The researcher is not interested in the donor of the
cells.

## The AI assistant's responses
The AI assistant's task is to help the researcher learn about the sample's biological nature. The
AI assistant has access only to a snapshot of the biological state of cellular sample
(represented via the provided information) and thereby can only respond with information that can
logically be derived from such a logical snapshot (e.g. no specific dosage numbers of treatments,
unimportant processing steps are likely to be reflected in such a snapshot, nor any donor- or
patient-specific information such as age or sex). Likewise, the AI assistant does not respond by
enumerating the provided genes and gene sets, but rather provides a more conceptual and
summarized form of the provided data, that describes the overall nature of the sample.
Furthermore, the AI assistant uses a tone of reflection and reasoning and without absolute
certainty (e.g. "It seems that ..."), and, where applicable, the AI assistant should include
reasoning steps to make the content more convincing and well-organized.
The responses must be fully in accordance with the provided information about the sample. If the
researcher asks a question that cannot be answered with the provided information, the AI
assistant should indicate so.

## Additional instructions
- refer to the data as "this sample" or "this cell type" or "these cells"
- keep it simple. no need to cover all information
- do not include any identifiers such as sample names, cell line names, ontology identifiers,
etc.
- Avoid the term 'gene sets'

## Format
Generate the conversation in the JSON format, following this schema:

[{"from": "researcher", "value": <str: the researcher's question>}, {"from": "ai", "value": <str:
the AI assistant's response>}, ...]
```

*Generated output for the "conversation" conversation prompt and the question-answer input for SRX7552910*

*Researcher:* What type of tissue does this sample represent?

*AI assistant:* This sample originates from renal tissue, specifically from a renal cell carcinoma tumor.

*Researcher:* What can you tell me about the state of this renal tissue?

*AI assistant:* Based on the provided information, the renal tissue appears to be cancerous, specifically classified as renal cell carcinoma at stage 2 without metastasis.

*Researcher:* What genes are highly expressed in this renal tissue?

*AI assistant:* The top expressed genes in this renal tissue include NEB, MUC12, C4A, IGHG1, IGHG1, SCARNA10, SNORD3D, IGLC1, IGFN1, CTAGE9, IGHG2, UGT2B7, SCARNA2, IGLC2, VWF, IGHGP, ENSG00000243302, USP17L15, NAIPP4, FLG, COL3A1, IGLC3, SCARNA7, PKHD1, ENSG00000284454, HRNR, LRRC37A, ARHGEF34P, COL1A2, ENSG00000281380, NAIPP2, ARHGEF35, IGKC, MGAM, RGPD1, AHNAK2, NPIPB12, PREX2, PPIAP22, LRRC37A3, ENSG00000215156, SST, SCARNA5, DNAH11, RGPD3, ENSG00000283738, USP17L6P, C3.

### 4. Generation of cluster labels for user-provided datasets

*Prompt for the generation of cell cluster descriptions using CellWhisperer's multimodal LLM*

User:

```
Help me analyzing this sample of cells. Always respond in proper english sentences and in a tone of uncertainty.
```

Assistant:

```
Sure thing. What do you want to know?
```

User:

```
Describe the biological state of these cells
```

*Prompt for condensation of the generated cluster descriptions using GPT-4*

```
Condense this information into a short title of maximum 8 words. Focus on the biological state, rather than the source or any specific perturbations of the sample
```
