## Supplementary Note 2 for "Multimodal learning of transcriptomes and text enables interactive single-cell RNA-seq data exploration with natural-language chats"

### Supplementary Note 2: Ablation study of the CellWhisperer multimodal model

The CellWhisperer multimodal AI model is based on various machine learning and data processing techniques. Here we analyze the importance of the individual techniques and model components, by training and testing adapted versions of our models that lacked one component at a time, in a process referred to as ablation (**Extended Data Fig. 1b**). We assessed the performance of these models on five representative metrics introduced in **Fig. 2**. For comparability and computational efficiency, all models were trained in four epochs with three different seeds. Below we describe each tested ablation and summarize the observed effects on performance.

First, CellWhisperer's multimodal embedding model follows the LiT method (Zhai et al. 2021), by initializing the network architecture and weights of the modality-specific embedding models based on pretrained transcriptome (Geneformer) and text (BioBERT) models, while freezing the former model during training. We tested the impact of different freezing configurations on the model performance. We observed that freezing both models or unfreezing only the transcriptome model led to a significant loss of performance, whereas freezing only the transcriptome model translated to the best performance among the tested configurations.

Second, we tested the use of an alterative transcriptome model as part of the CellWhisperer multimodal model. The scGPT model has achieved slightly better performance compared to Geneformer on various single-cell interpretation tasks (Cui et al. 2024; Kedzierska et al. 2023). We built an scGPT-based version of CellWhisperer and indeed observed increased performance on cell type prediction tasks. In contrast, the Geneformer-based version performed better in zero-shot disease prediction, indicating better ability to retain diverse biological information (beyond cell types) in the transcriptome embedding for diverse prediction tasks.

Third, we explored the effect of replacing the pretrained transcriptome model with a feed-forward neural network operating on the fixed-order gene expression vector and trained from scratch. As expected, results were substantially worse for the simpler model, but even that model retained better-than-random performance.

Fourth, we investigated the effect of training batch size on model performance, as it has previously been shown that contrastive training relies on particularly large batch sizes (Chen et al. 2020). We observed a significant decrease in performance when reducing the batch sizes by a factor of 16 (from 512 to 32), even when the gradients were accumulated across bags of 16 consecutive batches. We did not observe substantially improved results with larger batch sizes, nor with an alternative contrastive loss formulation (Jensen-Shannon divergence loss), which was described to perform well in low data and low batch size regimes (Shrivastava et al. 2023).

Fifth, we assessed the performance of multimodal models that were trained solely on data from one of the two data repositories (GEO or CELLxGENE Census). These models retained strong performance for most metrics, indicating that both repositories contain sufficiently broad information to support tasks such as zero-shot cell type prediction. Nevertheless, excluding the GEO dataset led to substantially reduced performance for disease prediction, which is likely due to the low representation of disease samples in CELLxGENE Census.

Sixth, we tested whether training data augmentation improves model performance. To that end, we generated slight variations of the textual annotations (based on different prompts in the LLM-assisted curation of the training data) and added individual single-cell profiles on top of the pseudo-bulk transcriptome profiles. Previous work on image-based CLIP models supports the value of data augmentation (Fan et al. 2023), but for CellWhisperer we did not observe a clear performance increase, hence we did not further pursue this strategy.

Finally, given the inherent biases of our training data (largely due to the unequal attention that different areas of biology have received and the resulting biases in the RNA-seq data submitted to GEO and other databases), we systematically increased the training weights of samples in areas with low dataset coverage, similarly to recent work on counter-balancing dataset biases in contrastive learning (Alabdulmohsin et al. 2024). However, we observed only negligible differences compared to unweighted training, indicating that different sample densities in our training data did not significantly affect model performance, as assessable through our metrics.

In summary, these analyses highlight the performance contributions of different aspects of our model architecture and training strategies, providing insights into the design of multimodal transcriptome-text models.
